# Synaptotagmin 7 outperforms synaptotagmin 1 to promote the formation of large, stable fusion pores via robust membrane penetration

**DOI:** 10.1101/2022.07.07.499215

**Authors:** Kevin C. Courtney, Taraknath Mandal, Nikunj Mehta, Lanxi Wu, Yueqi Li, Debasis Das, Qiang Cui, Edwin R. Chapman

**Affiliations:** Howard Hughes Medical Institute and the Department of Neuroscience, University of Wisconsin, 1111 Highland Avenue, Madison, Wisconsin, 53705; Department of Biochemistry and Molecular Medicine, West Virginia University, Morgantown, WV, 26506; Department of Chemistry, Boston University, Boston, MA, 02215; Department of Physics, Indian Institute of Technology – Kanpur, Kanpur 208016, India; Center for Bioanalytical Chemistry, University of Science and Technology of China, Hefei, 230026, China; Department of Biological Sciences, Tata Institute of Fundamental Research, Homi Bhabha Road, Navy Nagar, Colaba, Mumbai - 400005, India

## Abstract

Synaptotagmin-1 and synaptotagmin-7 are two prominent Ca^2+^ sensors that regulate exocytosis in neuronal and neuroendocrine cells. Upon binding Ca^2+^, both proteins partially penetrate lipid bilayers that bear anionic phospholipids, but the specific underlying mechanisms that enable them to trigger exocytosis remain controversial. Here, we examined the biophysical properties of these two synaptotagmin isoforms and compared their interactions with phospholipid membranes. We discovered that synaptotagmin-1•membrane interactions are greatly influenced by membrane order; tight packing of phosphatidylserine inhibits binding due to impaired membrane penetration. In contrast, synaptotagmin-7 exhibits robust membrane binding and penetration activity regardless of phospholipid acyl chain structure. Thus, synaptotagmin-7 is a “super-penetrator”. We exploited these observations to specifically isolate and examine the role of membrane penetration in synaptotagmin function. Using nanodisc-black lipid membrane electrophysiology, we demonstrate that membrane penetration is a critical component that underlies how synaptotagmin proteins regulate reconstituted, exocytic fusion pores in response to Ca^2+^.

## Introduction

Within the seventeen-member family of mammalian synaptotagmin (syt) isoforms^1^, syt1 and syt7 are currently the most heavily studied^2, 3^. It is now established that both isoforms play important roles in the synaptic vesicle (SV) cycle and neurotransmission, but their detailed mechanisms of action remain unclear. Both isoforms bind Ca^2+^ ions via each of their tandem C2-domains, designated C2A and C2B. Syt1 binds Ca^2+^ with lower affinity^4^, and responds to changes in [Ca^2+^] with faster kinetics than syt7^5^. Syt1 is localized to SVs^2, 6–8^, where it clamps spontaneous release^9, 10^ under resting conditions. Then, upon depolarization and Ca^2+^ entry^11, 12^, syt1 functions to trigger and synchronize evoked release^9, 13–15^. Syt1 also has additional functions, including the formation of the readily releasable pool of SVs^16^, SV docking^17^, and accelerating the kinetics of endocytosis^18, 19^. In contrast, syt7 largely resides on the axonal plasma membrane^20, 21^ in nerve terminals where it supports asynchronous neurotransmitter release^22, 23^. Additionally, syt7 KOs exhibit alterations in short term synaptic plasticity, including a loss of paired pulse facilitation ^24^ and enhanced synaptic depression with impaired SV replenishment^25^, without affecting spontaneous or single-stimulus evoked neurotransmitter release^25, 26^. Recently, syt7 was shown to promote activity-dependent docking of SVs at active zones^21, 23^, which may provide a unifying mechanism to support the multitude of functions described for this isoform. Notably, this syt7 docking function was determined to be upstream of Doc2α, a Ca^2+^ sensor for asynchronous neurotransmitter release^23, 27^. In addition to being targeted to the plasma membrane of axons, syt7 also resides on the surface of lysosomes^28^ to promote plasma membrane repair^29^. Furthermore, both syt1 and syt7 are present on dense core vesicles in neurons and in chromaffin cells where they are thought to serve as Ca^2+^ sensors for exocytosis^30, 31^. In addition, in chromaffin cells, syt7 has also been proposed to act as a docking/priming protein^32^. Whether syt7 is targeted to secretory vesicles or the plasma membrane might confer unique functional roles in membrane trafficking and exocytosis.

Although there is a long history of *in vitro* reconstitution and functional characterization of syt1^33^, to date, the successful purification of full-length syt7 has not been reported. This has precluded direct comparisons between full-length versions of these two isoforms in reduced systems. With respect to their molecular mechanisms of action, there is a consensus that upon binding Ca^2+^, the C2-domains syt1 and syt7 partially penetrate membranes that harbor anionic phospholipids^34, 35^. This membrane penetration step might serve to destabilize the local phospholipid environment by introducing volume into the bilayer, buckling the membrane, and lowering the energy barrier for fusion^34, 36–39^. Additionally, membrane penetration could enable syts to regulate SNARE complex assembly^40, 41^ and stabilize curved intermediate structures^42^. Finally, since syt1 and syt7 are membrane-anchored proteins, interactions between their C2-domains and ‘target’ membranes also serve to closely juxtapose the bilayers that are destined to merge, thus facilitating SNARE-mediated fusion^43^.

The importance of syt1 membrane penetration is supported by *in vitro* mutagenesis studies. In each C2-domain, the distal tips of two Ca^2+^ binding loops insert into membranes upon Ca^2+^ binding^33, 34, 43^. Substitution of residues at the tips of these loops, with four tryptophan residues (4W) to increase hydrophobicity and interfacial interactions with phospholipids, enhances membrane binding and bending, while substitution with alanines (4A) reduces membrane association^39, 44^. Moreover, when these mutant proteins are expressed in syt1 KO neurons, 4A fails to rescue synchronous neurotransmitter release whereas 4W expression exhibits a significant increase in EPSC amplitude and Ca^2+^-sensitivity, compared to WT syt1^45, 46^. However, the 4W and 4A mutations have other effects on syt1 biochemistry, including altered interactions with SNARE proteins^39^. Additional approaches are needed to unambiguously define the precise role of syt1•membrane interactions in fusion.

Here, we examine how membrane binding and penetration by the C2-domains of syt1 and syt7 contribute to Ca^2+^-triggered membrane fusion. Specifically, we use phospholipid bilayer order as a tool to explore the link between Ca^2+^•syt membrane penetration and the regulation of reconstituted fusion pores, which represent the first crucial intermediate in the fusion reaction. We demonstrate that syt function can be controlled by manipulating phosphatidylserine (PS) acyl chain structure. Namely, syt1 did not bind or penetrate bilayers that contained PS with saturated acyl chains; without efficient membrane penetration, syt1 failed to promote or stabilize fusion pores in response to Ca^2+^. Surprisingly, we observed that syt7 displays far more robust membrane penetration activity as compared to syt1, and efficiently penetrated all bilayers that were tested, regardless of the membrane order. Furthermore, we report the first use and characterization of reconstituted full-length syt7 and determine, via comparisons with syt1, that the ability of their C2-domains to penetrate membranes is a crucial step in the regulation of fusion pores.

## Results

### Syt7, but not syt1, penetrates membranes that harbor saturated PS

To compare how syt1 and syt7 interact with phospholipid bilayers, we first performed a well-described syt•loop penetration assay^34, 43^. For this, the cytoplasmic domains (denoted C2AB) of each isoform were labeled with a solvatochromic fluorescent dye, NBD, on a cysteine residue placed in the distal tip of a membrane penetration loop (loop-3); we labeled the C2B of syt1, the dominant C2-domain^47, 48^, and either C2A or C2B of syt7. In the absence of Ca^2+^ (+EGTA), these C2AB domains only weakly interact with 100 nm PC/PS (80:20) liposomes, resulting in low fluorescence signals (**Figs. 1a-c and Supplementary Fig. 2**). Upon binding Ca^2+^, the C2AB domains rapidly associate with liposomes and partially insert into the hydrophobic core of the bilayer^34, 35^, causing a significant increase, and a blue-shift, in NBD fluorescence (**Figs. 1a-c and Supplementary Fig. 2**). To specifically examine the contribution of hydrophobic interactions to the binding reaction, we compared the ability of syt1 and syt7 to penetrate bilayers containing 20% PS with either saturated (16:0/16:0) or unsaturated acyl chains (18:1/18:1), also notated as dipalmitate-phosphatidylserine (DPPS) and dioleate-phosphatidylserine (DOPS), respectively; the remaining 80% of the lipids in both conditions were unsaturated dioleate-phosphatidylcholine (DOPC) (**Supplementary Fig. 1)**. Knowing that syt1 and syt7 are PS binding proteins, we postulated that tight lateral packing of saturated PS acyl chains in the liposomes might render the bilayers refractory to penetration, thus restricting hydrophobic interactions. We first confirmed that syt1 and syt7 efficiently penetrate bilayers containing unsaturated DOPS (**Figs. 1b & 1c and Supplementary Fig. 2**). However, if the PS acyl chains are fully saturated, syt1 penetration was virtually abolished (**Fig. 1b**), despite having 80% of the bilayer composed of unsaturated DOPC. Surprisingly, we found the C2A and C2B domains of syt7 C2AB exhibited comparable penetration into saturated and unsaturated PS bilayers (**Fig. 1c and Supplementary Fig. 2**), indicating a more robust mode of interacting with membranes.

**Figure 1.**
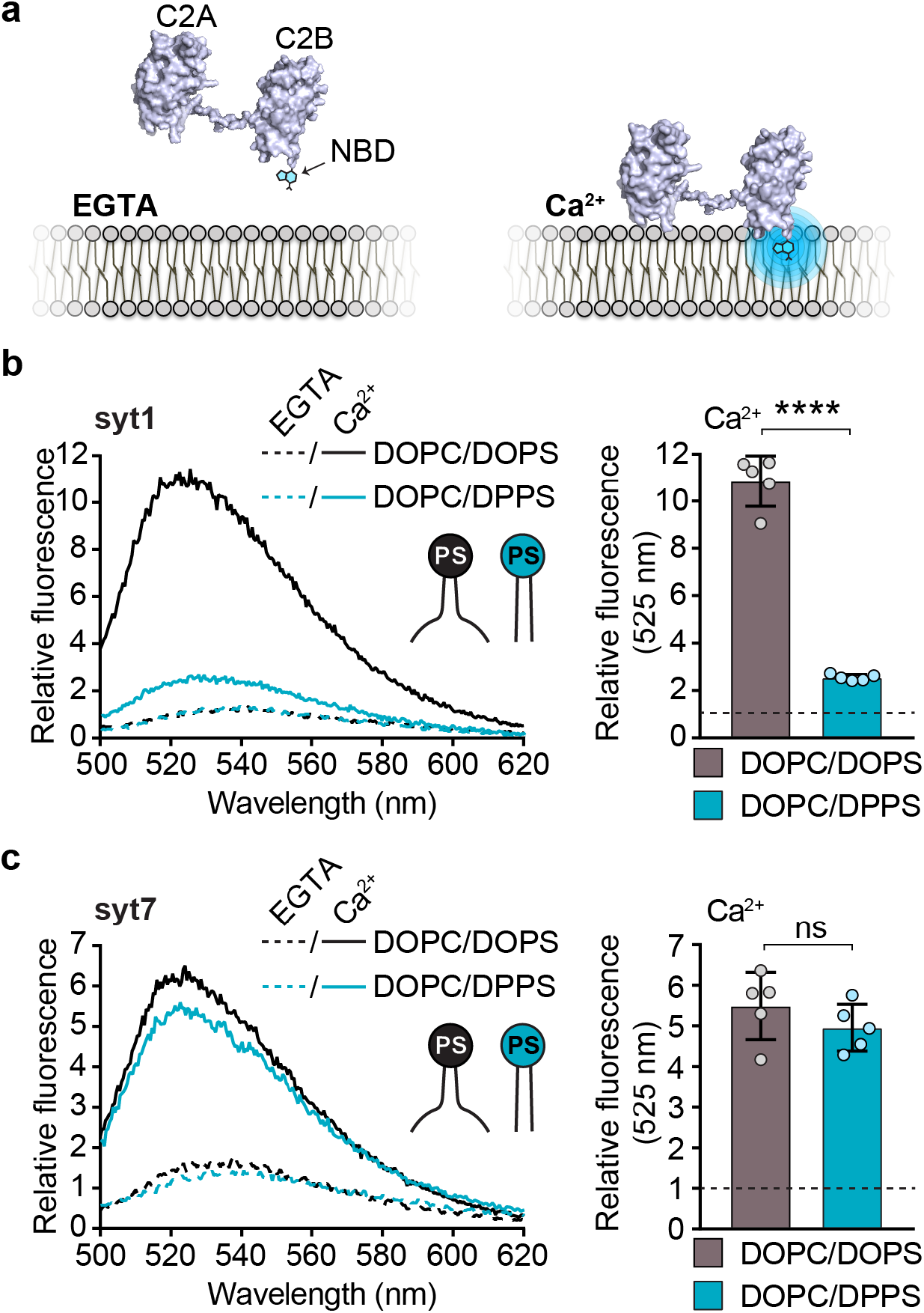
Syt7, but not syt1, penetrates membranes that harbor saturated PS. **a** Illustration showing how an NBD labelled C2AB domain associates with a lipid bilayer in response to binding Ca^2+^. After binding Ca^2+^, the C2AB domain binds and partially penetrates the membrane, thus inserting the NBD dye into the hydrophobic core of the bilayer, causing a blue shift and an increase in fluorescence intensity. **b** Representative fluorescence emission spectra (*left panel*) of NBD labelled syt1, in the presence (solid lines) and absence (dotted lines) of Ca^2+^, and liposomes composed of 80:20 DOPC/DOPS (black) or DOPC/DPPS (blue). The syt1 C2AB domain is labeled on loop 3 of the C2B domain at position 367. Quantification of NBD-syt1 C2AB fluorescence emission at 525 nm in the presence of Ca^2+^ and liposomes composed of DOPC/DOPS (black) or DOPC/DPPS (blue), *right panel*. The data are normalized to the EGTA condition, shown as a horizontal black dotted line. **c** Representative fluorescence emission spectra (*left panel*) and quantification (*right panel*) of NBD labelled syt7, under the same conditions as *panel b*. The syt7 C2AB domain is labeled on loop 3 of the C2B domain at position 361. Each condition was repeated five times on different days using fresh materials. Error bars represent standard error of the mean. **** represents p<0.0001 and ns represents a non-significant difference between conditions determined by Student’s t-test.

To establish the generality of the syt1 penetration defect observed using saturated PS-bilayers, we examined the membrane penetration performance of other C2-domain-containing proteins under the same conditions as were used with syt1 and syt7. We engineered a single cysteine substitution into the third loop of the C2-domains from protein kinase C (PKC) and cytosolic phospholipase A2 (cPLA2), as well as the third loop in the C2B domain of Doc2β C2AB. We found that all three proteins also displayed significantly impaired penetration into membranes with saturated acyl chains in response to Ca^2+^ (**Supplementary Fig. 3**), thus the robust penetration of syt7 into DPPS bilayers appears unique.

### Molecular dynamics simulations show saturated PS clusters resist syt1 C2B-domain penetration

Next, to further examine how syt1 and syt7 interact with bilayers containing saturated and unsaturated PS, we conducted all-atom molecular dynamics (MD) simulations of their C2-domains with phospholipid bilayers, again using the same lipid compositions used above. We first performed lipid-only MD simulations to monitor the stability of DOPS and DPPS clusters in the DOPC/DxPS (80:20) bilayers. We postulated that DPPS may form a tight cluster, thus restricting C2-domain penetration. For this initial test, a small cluster of 32 PS lipids, either DOPS or DPPS, was placed in the center of a DOPC membrane (**Supplementary Fig. 4a**) and equilibrated for 450 ns using MD simulations. **Fig. 2a** shows that DPPS lipids (cyan) are much more compact than DOPS lipids (blue), suggesting that the saturated PS lipids are likely to form bigger, more stable clusters in a saturated-unsaturated lipid mixture (**Supplementary movies 1 & 2**). We then quantified how clustered the two PS species were at the end of the simulations by measuring the midpoint distances between the PS molecules; the radial distribution function (RDF), shows that the DPPS molecules remain in close proximity with a peak centered around 0.75 nm. In contrast, the DOPS molecules are more dispersed, as indicated by a shorter peak height with a broad distribution (**Fig. 2b**). These initial MD simulations were also validated via atomic force microscopy (AFM) imaging of supported lipid bilayers containing 20% DOPS or DPPS; we found that Ca^2+^ reversibly caused DPPS to segregate into 15 ± 7 nm clusters in the bilayer (**Supplementary Fig. 5a**). In contrast, no lipid clustering was observed in the DOPS condition (**Supplementary Fig. 5b**).

**Figure 2.**
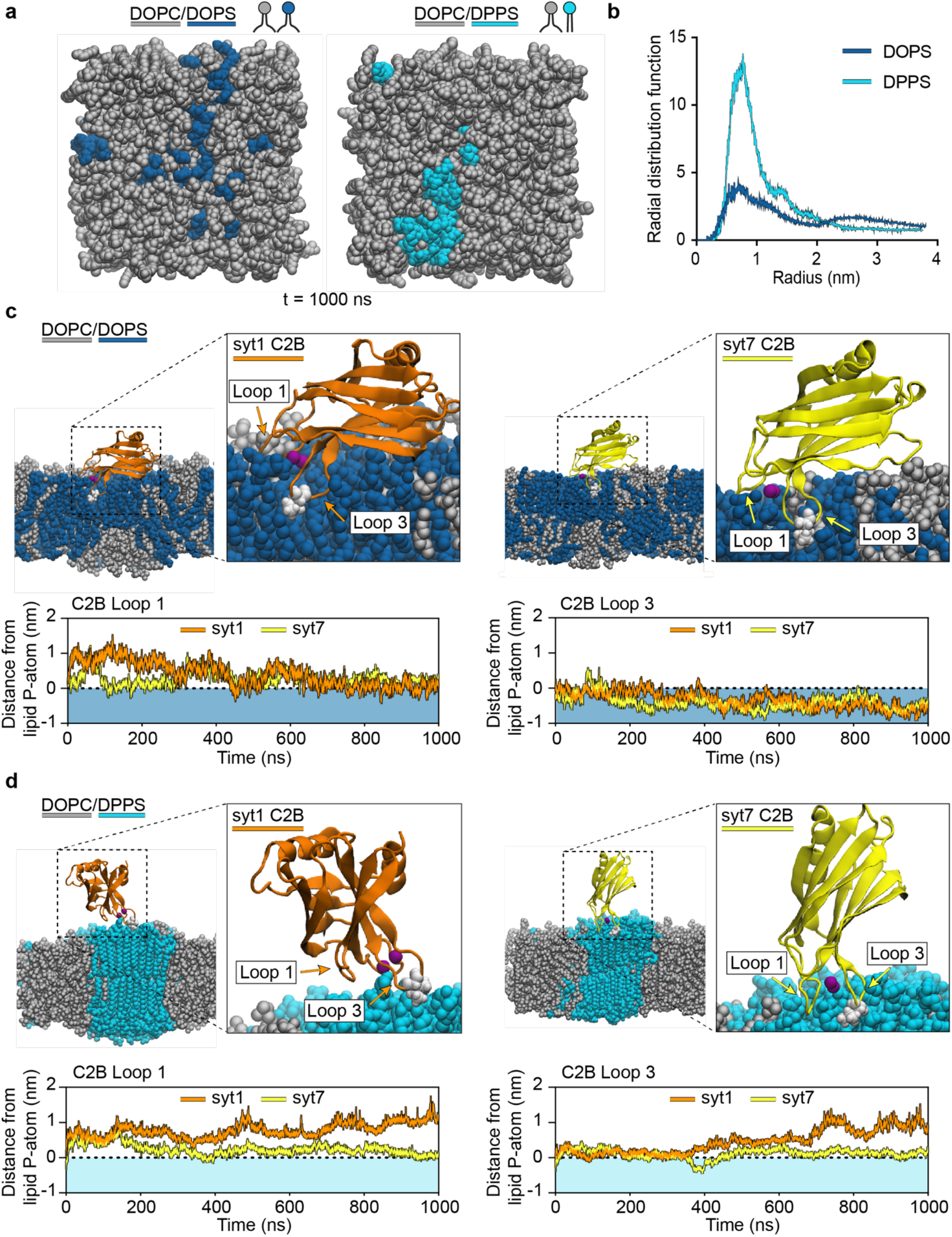
Molecular dynamics simulations of syt1 and syt7 C2B•membrane interactions. **a** End-point (450 ns) MD simulations of lipid bilayers composed of DOPC/DOPS or DOPC/DPPS. The PS lipids were initially placed as a cluster in the center of the membrane (see Supplementary Fig. 4 for the starting images) and then allowed to freely diffuse over time. **b** Radial distribution functions comparing the clustering behavior of DOPS and DPPS in the lipid bilayer at the end of the simulation shown in *panel a*. **c** End point (1000 ns) MD simulations snapshots showing syt1 (orange) and syt7 (yellow) C2B domains interacting with lipid bilayers composed of DOPC/DOPS. Quantification of loop 1 and loop 3 depth from syt1 and syt7 into the DOPS-containing bilayer (blue shading) are shown in the lower panels. In each case, the loop depth is normalized relative to the position of the lipid phosphate group, indicated by a horizontal dotted line. **d** End point (1000 ns) MD simulations snapshots showing syt1 (orange) and syt7 (yellow) C2B domains interacting with lipid bilayers composed of DOPC/DPPS. Quantification of loop 1 and loop 3 depth from syt1 and syt7 into the DPPS-containing bilayer (light cyan shading) are shown in the lower panels. The loop residue that achieved the deepest depth of penetration, I367 for syt1 and L361 for syt7, is emphasized in white.

We then carried out a second set of MD simulations to examine how PS acyl chain structure affects C2-domain•membrane interactions using syt1 C2B, syt7 C2A, and syt7 C2B. Each isolated C2B domain was placed on top of the membrane such that the membrane penetration loops were positioned close to the PS cluster (**Supplementary Figs. 4b & 4c**). The system was solvated and charge neutralized by adding Ca^2+^ and Cl^-^ ions. Upon equilibration of the C2-domains, we found that Ca^2+^-bound loops 1 and 3 of the C2B domains spontaneously inserted into, and remained inside, the DOPC/DOPS membrane throughout the 1000 ns long MD simulations (**Fig. 2c**). In the case of syt7 C2A, however, tilting of the domain in favor of loop 3 penetration restricted loop 1 access to the hydrophobic core of the bilayer (**Supplementary Fig. 6a**). In all cases using DOPS, the insertion depth of loop 3 was greater than the depth of loop 1, while loop 2 remained away from the lipid head groups (**Fig. 2c**, *lower panel* **and Supplementary Fig. 6a**). Insertion of loops 1 and 3 allows the hydrophobic residues, V304 and I367 of syt1 C2B, F229 of syt7 C2A, and I298 and L361 of syt7 C2B, to access the hydrophobic bilayer core, strengthening the binding interaction. This result matches well with previous experimental results from our laboratory^43^. In contrast to the DOPS simulations, tight packing of the PS lipids in the DOPC/DPPS membrane does not allow strong binding or insertion of the syt1 C2B domain (**Fig. 2d**, *left panel*). However, in line with our experimental results (**Fig. 1c and Supplementary Fig. 2**), we found that the syt7 C2A and C2B domain loops penetrated well into the DPPS cluster (**Fig. 2d**, *right panel* **and Supplementary Fig. 6**). Additional pairwise comparisons of the penetration depths that were reached by the C2-domain loops are shown in **Supplementary Fig. 7**.

### Cholesterol rescues the syt1 membrane penetration defect

The experiments and MD simulations above reveal that acyl chain packing strongly affects the ability of syt1, but not syt7, to penetrate membranes in response to Ca^2+^. We therefore tested whether ‘loosening’ the lateral packing of DPPS bilayers can rescue syt1 penetration. Physiological plasma membrane lipid bilayers contain approximately 30 - 40% cholesterol^6, 49, 50^. Cholesterol displays unique membrane modulating properties by reducing the rotational mobility (stiffening) of unsaturated acyl chains, while also adding space between (loosening) tightly packed saturated acyl chains. We therefore hypothesized that introducing cholesterol into DPPS bilayers would loosen the PS clusters and allow the syt1 C2B domain to penetrate otherwise refractory membranes. After repeating the DPPS MD simulations in the presence of 30% cholesterol, we indeed found that loops 1 and 3 of syt1 C2B could penetrate the DPPS membrane (**Figs. 3a & 3b**). Correspondingly, in AFM experiments, Ca^2+^ failed to induce detectable clusters in DPPS supported lipid bilayers that contained 30% cholesterol (**Supplementary Fig. 5c**).

**Figure 3.**
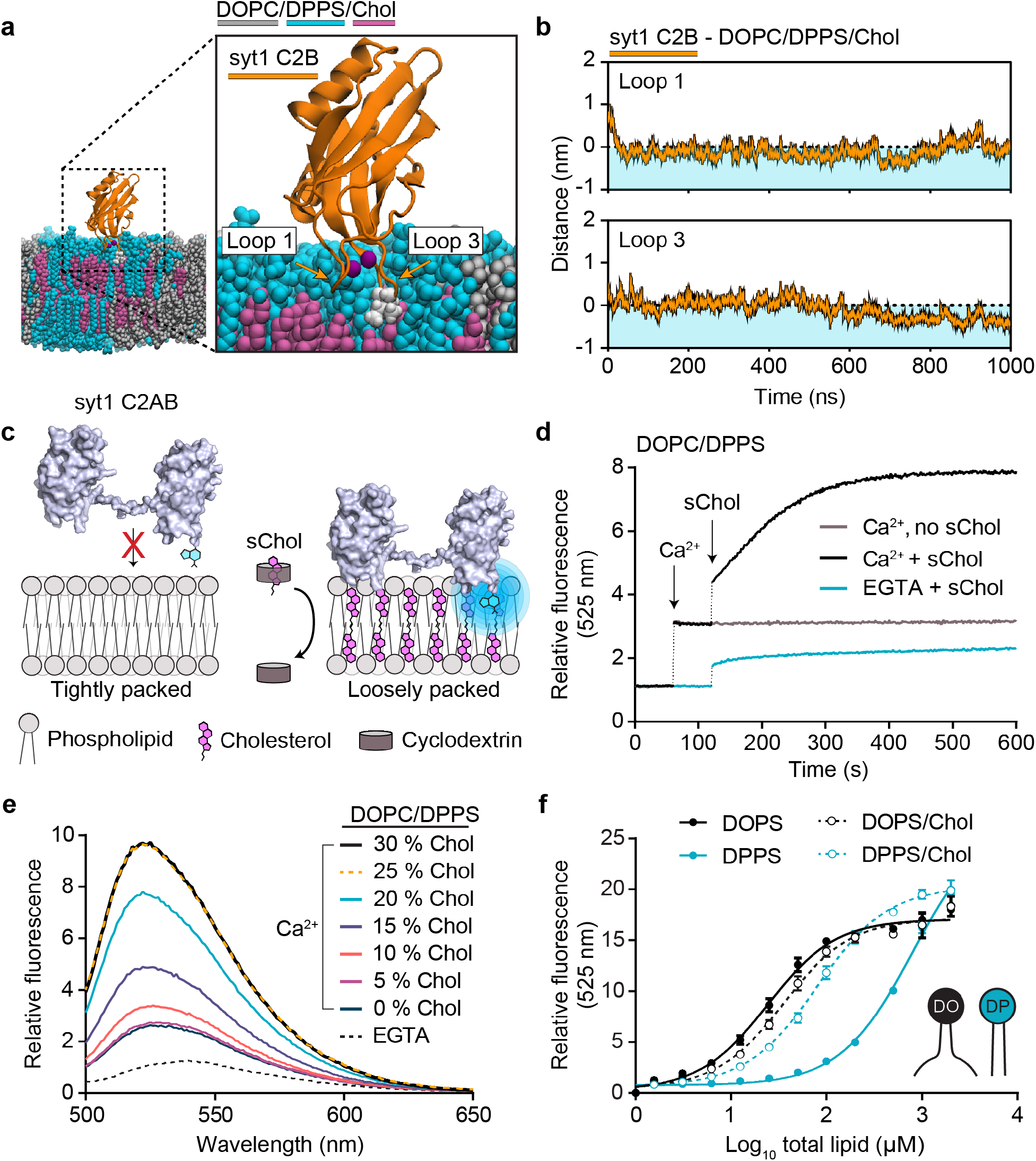
Cholesterol enables syt1 to penetrate bilayers containing DPPS. **a** MD simulations snapshot after 1000ns of a syt1 C2B domain (shown in orange) interacting with a phospholipid bilayer composed of DOPC/DPPS/cholesterol. DOPC is shown in grey, DPPS in cyan and cholesterol shown in magenta. Residue I367 on loop 3 of syt1 is emphasized in white and the bound Ca^2+^ ions are shown in purple. **b** Quantification of the penetration depth of loop 1 (*top panel*) and loop 3 (*lower panel*) from syt1 C2B domain into the DPPS-containing bilayer (light cyan shading) throughout the 1000 ns MD simulation. The loop depth is normalized relative to the position of the lipid phosphate group, indicated by a horizontal dotted line. **c** An illustration depicting experimental C2 domain membrane penetration assay with the addition of a cholesterol-cyclodextrin complex, known as soluble cholesterol (sChol). In the absence of cholesterol, loop 3 in the C2B domain of syt1 C2AB only minimally penetrates the bilayer containing DPPS. When sChol is applied, cholesterol is donated into the DPPS bilayer, which enables syt1 C2AB to efficiently penetrate the membrane. **d** Representative time course examining NBD-labelled syt1 C2AB fluorescence in the presence of DOPC/DPPS (80:20) under the indicated conditions. Note: the fluorometer was briefly paused during the addition and mixing of Ca^2+^ and sChol into the cuvette, indicated by the vertical dashed lines. **e** Representative fluorescence spectra of NBD-labelled syt1 C2AB with various 100 nm liposome populations composed of DOPC/DPPS and increasing cholesterol. **f** Quantification of a liposome titration in the presence of NBD-labelled syt1 C2AB (I367C). The experiments were repeated one three separate occasions with fresh materials. The protein concentration was fixed at 250 nM and the fluorescence at 525 nm was monitored as the concentration of lipid increased from 0-500 µM. Error bars represent standard error of the mean from triplicate experiments performed on separate days with fresh materials.

Next, we revisited the syt1 NBD•loop penetration experiments described in **Fig. 1** to further examine the effects of cholesterol, experimentally. First, we used soluble cholesterol-loaded methyl-β-cyclodextrin (sChol) to introduce cholesterol into DOPC/DPPS bilayers while monitoring changes in NBD fluorescence as a function of time (**Fig. 3c**). As described above (**Fig. 1b**), Ca^2+^-bound syt1 C2AB can only minimally penetrate DPPS-containing bilayers, as indicated by a relatively low fluorescence signal under this condition (**Fig. 3d**, *grey trace*). However, after of the addition of sChol, to incorporate cholesterol into the liposomes, syt1 C2AB regained the ability to penetrate the bilayer (**Fig. 3d**, *black trace*), showing maximal penetration after approximately 10 minutes.

We then generated 100 nm DOPC/DPPS liposomes with increasing amounts of cholesterol, up to 30%, while monitoring membrane penetration by fluorometry. As with the real-time addition of cholesterol, discrete increases in the cholesterol concentration allowed syt1 C2AB to penetrate membranes, with an EC_50_ of 17.6%, reaching maximal membrane penetration at 25% cholesterol (**Fig. 3e**). Without cholesterol, approximately ten-fold more lipid was needed to drive maximal penetration of syt1 into liposomes bearing DPPS (protein:lipid, 1:4000), compared to DOPS (protein:lipid, 1:400) (**Fig. 3f**). However, if cholesterol is present, syt1 penetration is comparable for all PS species, across a wide range of protein/lipid ratios (**Fig. 3f**). This effect of cholesterol on syt1 binding to DPPS rationalizes how Kiessling *et al.* (2018) observed syt1 triggering membrane fusion using bilayers containing saturated PS, as these membranes also contained 20% cholesterol^51^. Together, these results support the conclusion that syt1 fails to penetrate DPPS-containing membranes, due to the rigidity of the bilayer. However, the attributes of syt7 that enabled robust penetration into DPPS bilayers remained unclear; this is addressed further below.

### C2-domain binding to phospholipid bilayers is coupled to membrane penetration

To determine whether the compromised penetration of syt1 into DPPS-containing bilayers was also accompanied by a failure to bind to the liposome surface via electrostatic interactions, we performed a protein-liposome co-sedimentation assay. The C2AB domains were mixed with 100 nm liposomes that were composed of 20% saturated (DPPS) or unsaturated PS (DOPS), in 0.2 mM EGTA or 0.5 mM free Ca^2+^, followed by ultracentrifugation to pellet the liposomes (**Fig. 4a**); bound proteins co-sediment with the liposomes. C2AB domains from both syt1 and syt7 efficiently bound DOPS-containing liposomes in the presence of Ca^2+^, as indicated by a depletion of protein from the supernatant (**Fig. 4b**). However, analogous to the penetration data, only 20-25% of syt1 C2AB bound DPPS-bearing liposomes in Ca^2+^ (**Fig. 4c**). This validates that membrane insertion is a prerequisite for efficient syt1•membrane binding activity, presumably by providing hydrophobic interactions; electrostatic interactions alone are apparently insufficient to enable syt1 to stably bind anionic lipid membranes. This conclusion is consistent with earlier mutagenesis studies^39^. In contrast to syt1, and in agreement with the penetration results, we found that syt7 maintained robust binding to liposomes bearing either unsaturated or saturated PS (**Figs. 4b & 4c**), further supporting the idea that membrane binding is coupled with penetration. Notably, we consistently observed that approximately 25-30% of the syt7 bound to the liposomes in EGTA; this might reflect the propensity of DPPS to form clusters, thereby increasing the local concentration of this lipid. Indeed, a previous report showed that syt7 exhibits robust Ca^2+^-independent binding to liposomes, as a function of the mole fraction of PS^52^.

**Figure 4.**
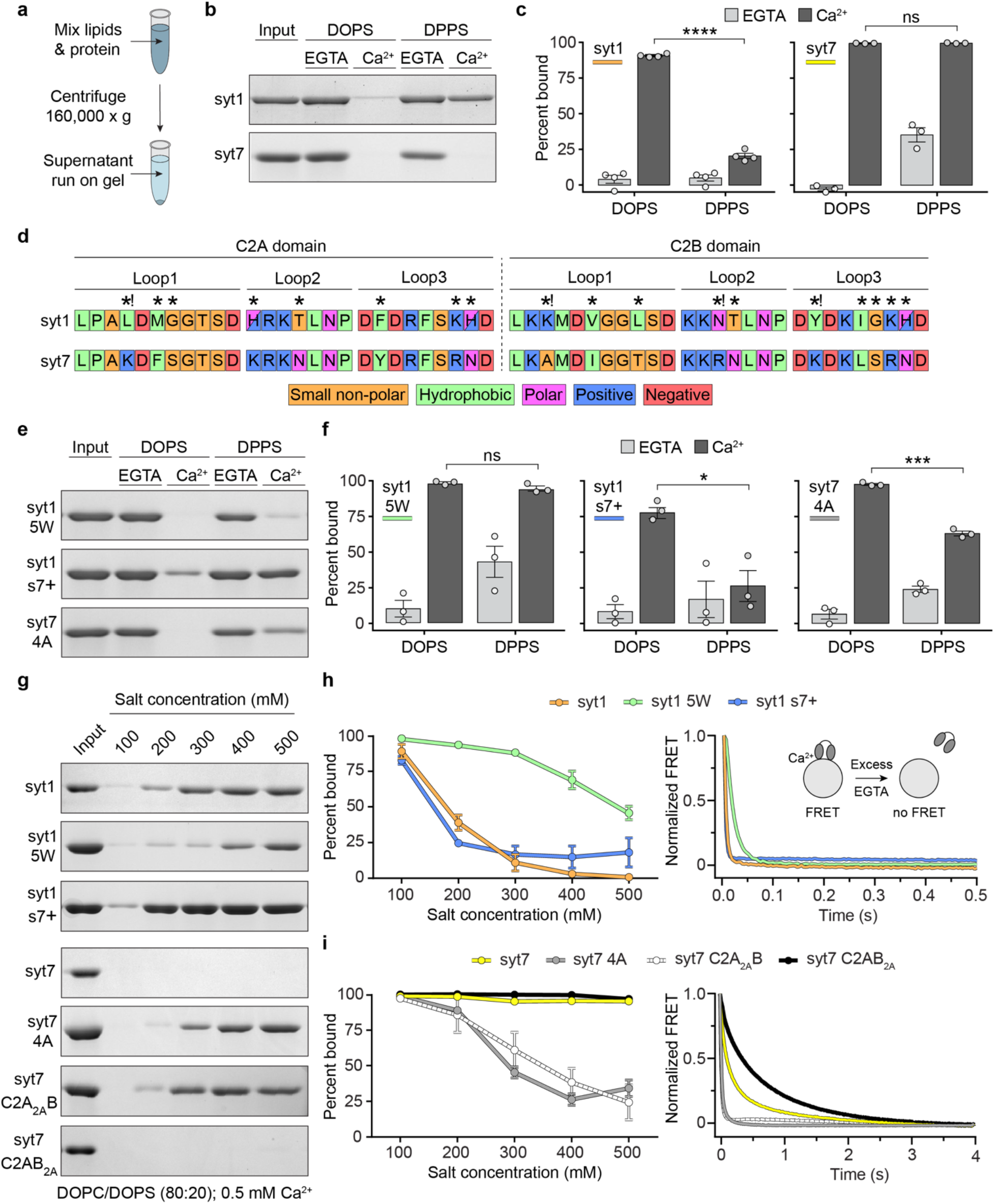
Hydrophobic interactions dominate syt1 and syt7 membrane association. **a** Illustration of the liposome-protein co-sedimentation assay. Proteins and liposomes are mixed, followed by sedimentation of the liposomes; the supernatant is subjected to SDS-PAGE to assay for protein depletion. **b** Representative Coomassie stained SDS-PAGE gels of syt1 and syt7 co-sedimentation samples, in the presence and absence of Ca^2+^, using liposomes composed of DOPC/DOPS or DOPC/DPPS. Throughout the figure, “Input” refers to the protein-only control sample. **c** Quantification of the replicated syt1 and syt7 co-sedimentation assays comparing binding to DOPS and DPPS containing liposomes. **d** Amino acid sequences of the penetration loops of syt1 and syt7 C2A and C2B domains. Unique residues at comparable positions between the two sequences are indicated by an asterisk (*). Unique residues with a differences in charge are indicated by asterisk and exclamation point (*!). **e** Representative Coomassie stained SDS-PAGE gels of mutant syt1 and syt7 co-sedimentation samples, in the presence and absence of Ca^2+^, and liposomes composed of DOPC/DOPS or DOPC/DPPS. **f** Quantification of the mutant syt1 and syt7 co-sedimentation assays comparing binding to DOPS- and DPPS-bearing liposomes. **g** Representative Coomassie stained SDS-PAGE gels of WT and mutant syt1 and syt7 co-sedimentation samples containing between 100 and 500 mM salt. **h** Quantification of the WT and mutant syt1 co-sedimentation samples containing between 100- and 500-mM salt (*left panel*). Disassembly of Ca^2+^-dependent WT and mutant syt1 complexes with liposomes, measured by stopped flow rapid mixing with the Ca^2+^ chelator, EGTA (*right panel*). **i** Quantification of experiments performed using WT and mutant syt7, as described in panel h. Each condition was repeated at least three times on different days using fresh materials. Error bars represent standard error of the mean. Conditions were compared using the Student’s t-test. Within the bar graphs, ****, ***, * and ns represents p<0.0001, p<0.001, p<0.5 and a non-significant difference, respectively.

### Hydrophobic residues are critical for C2-domain•membrane interactions

The membrane binding and penetration assays described above revealed that the C2-domains of syt1 and syt7 engage with lipid bilayers in somewhat distinct manners. We therefore compared the amino acid sequences of their membrane penetration loops; overall, the sequences are highly conserved (**Fig. 4d**). The most notable differences are that the syt1 loops contain three more hydrophobic residues, and three fewer positive charged residues (excluding the histidine residues in the syt1 loops at physiological pH) than the syt7 loops. This could be expected to equip syt1 with comparatively enhanced penetration activity, and syt7 with a prominent electrostatic binding component; instead, we found that syt7 displayed enhanced membrane binding (**Figs. 4b & 4c**) and penetration activity (**Fig. 1c**). To delve into this further, we generated a series of mutants to examine how the hydrophobic and cationic character of the loops influenced binding. These included a syt1 C2AB construct that contained five tryptophan substitutions in the penetration loops (syt1 5W) to increase hydrophobic and interfacial interactions with phospholipids, or a poly-cationic mutant (L171K, H198K, K236R, N333R, Y364K, K369R) that grafted all the cationic residues of the syt7 loops onto syt1 (syt1 s7+). Note that while K236R and K369R in the syt1 s7+ construct do not affect the charge at these two positions, arginine residues have been reported to exhibit enhanced interfacial binding to phospholipids, compared to lysines^53^. We also examined a syt7 C2AB mutant with reduced hydrophobicity (F167, F229, I298, L361A) in the penetration loops (syt7 4A).

We first tested how syt1 5W, syt1 s7+ and syt7 4A performed against the refractory bilayers containing saturated DPPS. Co-sedimentation assays revealed that syt1 5W exhibited enhanced membrane binding activity in response to Ca^2+^; the tryptophan substitutions enabled syt1 5W to bind equally well to DOPS and DPPS (**Figs. 4e & 4f**). Moreover, NBD-labelled syt1 5W was able to efficiently penetrate DPPS bilayers, revealing a clear gain-of-function (**Supplementary Fig. 8**). In contrast, the syt1 s7+ construct did not improve binding to DPPS in response to Ca^2+^ (**Figs. 4e & 4f**) and displayed a partial reduction in binding to DOPS in Ca^2+^ (**Figs. 4e & 4f**). In short, this mutant exhibited the opposite of the predicted effect. When examining syt7 4A, we found this construct had a reduced ability to bind DPPS in Ca^2+^, compared to WT syt7 (**Figs. 4c, 4e & 4f**), again highlighting the importance of hydrophobic residues in the loops.

To further explore how electrostatic interactions contribute to syt•membrane interactions, we conducted protein-liposome co-sedimentation experiments as a function of increasing ionic strength. We confirmed that syt1•membrane binding was highly sensitive to the salt concentration (**Figs. 4g & 4h,** *left panel*), whereas WT syt7 was entirely resistant (**Figs. 4g & 4i,** *left panel*), as reported previously^34, 43, 54^. We then repeated these experiments using syt1 5W, syt1 s7+ and syt7 4A. We found that the syt1 5W substitutions had a greater impact on overcoming salt sensitivity, compared with positive charged residue substitutions (**Figs. 4g & 4h,** *left panel*); grafting the charged residues in the loops of syt7 onto syt1 (syt1 s7+) did not improve salt resistance. The syt1 s7+ results are surprising considering this construct contains prominent hydrophobic and electrostatic character yet is still outperformed by WT syt7. Moreover, syt1 5W was also outperformed by WT syt7 in this assay. In line with the DPPS co-sedimentation experiments (**Figs. 4e & 4f**), we also observed that hydrophobicity is critical for salt-insensitivity of syt7, as the 4A mutation enabled modest increases in salt (200 mM) to disrupt binding (**Figs. 4g & 4i,** *left panel*). To tease apart contributions of each syt7 C2-domain, we generated and tested syt7 C2AB constructs with reduced loop hydrophobicity in either the C2A (syt7 C2A_2A_B) or C2B (syt7 C2AB_2A_) domain. These results demonstrated that the C2A domain of syt7 is responsible for the salt insensitivity, whereas the syt7 C2AB_2A_ mutation had no effect (**Figs. 4g & 4i,** *left panel*). We also found, via stopped-flow rapid mixing experiments, that the membrane dissociation kinetics of each of these constructs shared the same trends as the co-sedimentation studies (**Figs. 4h & i,** *right panels*). Specifically, the 5W mutation slowed the disassembly of Ca^2+^•syt1 from liposomes after mixing with excess EGTA (**Fig. 4h**, *right panel*), and both the syt7 4A and syt7 C2A_2A_B mutations dramatically increased the disassembly rate (**Fig. 4i**, *right panel*), further supporting the importance of loop hydrophobicity. Curiously, we found that the syt7 C2AB_2A_ mutation reproducibly slowed the kinetics of disassembly (**Fig. 4i**, *right panel*). At present, a mechanism for this perplexing C2AB_2A_ mutation effect is unknown.

Our data support previous conclusions^39, 45^ that C2-domain loop hydrophobicity is critical for membrane binding, especially when contrasting our syt7 4A and syt1 s7+ results (**Figs. 4e & 4f**). However, considering syt7 has higher membrane binding affinity (**Supplementary Fig. 9**), with fewer hydrophobic residues in the loops, compared to syt1 (**Fig. 4d**), these results highlight the somewhat counterintuitive complexity of C2-domain•membrane interactions, which cannot easily be predicted from the primary sequence. These results indicate that the hydrophobicity of the C2-domain loops may be more important than the cationic properties in governing membrane binding. However, additional factors, other than the sum of hydrophobic and electrostatic residues in the loops, appear to influence C2AB•membrane interactions.

### Membrane penetration is required for synaptotagmin 1 to promote the formation of large, stable, fusion pores

Having established that PS acyl chain order can govern how C2-domains interact with lipid bilayers, we proceeded to investigate the functional consequences of restricting syt membrane penetration. We reasoned that PS acyl chain structure could be exploited to specifically interrogate the relationship between syt•membrane penetration and the ability of syts to promote membrane fusion^41^. For this, we used the recently developed nanodisc-black lipid membrane (ND-BLM) planar lipid bilayer electrophysiology approach^41, 55^. This involves the reconstitution of v-SNARE proteins into a nanodisc (ND) to act as a SV mimic, while t-SNAREs are reconstituted into a planar lipid bilayer to mimic the presynaptic plasma membrane. When v-SNAREs in NDs associates with t-SNAREs in the supported bilayer, *trans*-SNARE complexes assemble to creates a nascent fusion pore across the two membranes (**Supplementary Fig. 10**). The properties of these fusion pores are then monitored electrophysiologically. Previously, we reported that 13 nm NDs with three copies of the SV SNARE, synaptobrevin 2, (ND_3_) make small unstable fusion pores^42, 55^ with BLMs containing the t-SNAREs syntaxin-1A and SNAP-25B. However, incorporation of syt1 into these NDs caused the fusion pores to expand into larger, more stable structures in response to Ca^2+^ binding^41^. This experimental system is, therefore, ideal to examine the direct impact of syts on the properties of fusion pores. By substituting PS species within the BLM, we can examine how syt membrane penetration influences pore kinetic properties and dilation.

Classically, BLM experiments are conducted by first dissolving a lipid mixture in a solvent, such as n-decane, followed by using a fine tipped brush or glass capillary to paint the lipids across a small aperture to form a planar lipid bilayer^41^. We found that the critical lipid in this study, DPPS, is insoluble in n-decane, making the standard BLM protocol unusable. We therefore developed a new strategy to form a planar lipid bilayer across the aperture of a standard BLM cup that could accommodate all lipids used in this study (described in detail in the Methods section). This approach involves resuspending the lipids in pure water and then pipetting a droplet of lipid across the aperture of the BLM cup (**Fig. 5a**). The cup is then placed into a vacuum desiccator to evaporate the water and form a lipid film. Buffer is then applied, and the lipids self-assemble to form a bilayer across the aperture. In contrast to other BLM protocols, this new strategy also has the advantage of allowing the incorporation of t-SNAREs directly into the BLM lipid mixture, rather than requiring a second step to donate the t-SNAREs into the BLM after formation, as was done previously^41^. This drying method was successfully able to form t-SNARE-incorporated bilayers, enabling DPPS ND-BLM experiments.

**Figure 5.**
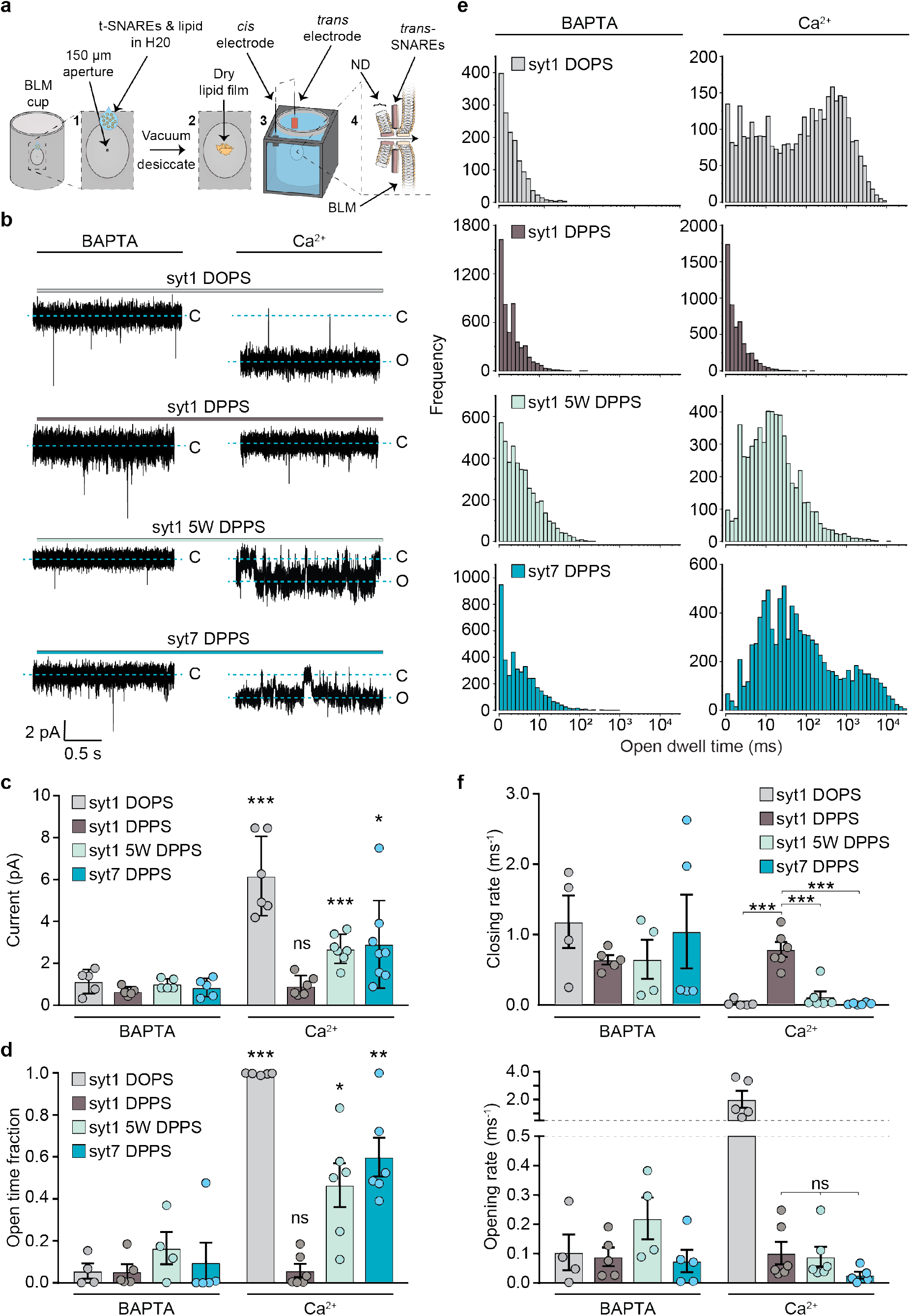
Membrane penetration is required for syts to trigger the dilated open state of fusion pores. **a** Illustration of the modified BLM protocol using lipid desiccation, which facilitated the formation of planar lipid bilayers containing DPPS. **b** Representative raw traces of syt1, syt1 5W and syt7 ND-BLM recordings. ND3-syt1 experiments were performed with 20% DOPS in the BLM as a positive control for the effect of Ca^2+^ on fusion pore properties The remaining traces were performed with 20% DPPS in the BLM, in 0.5 mM BAPTA or 0.5 mM free Ca^2+^. **c** Quantification of the current passing through ND-BLM fusion pores under the indicated conditions. The statistical notations refer to comparisons between the corresponding BAPTA and Ca^2+^ conditions. **d** Quantification of the fraction of time that ND-BLM fusion pores remained in the open state under the indicated conditions. The statistical notations refer to comparisons between the corresponding BAPTA and Ca^2+^ conditions. **e** Open dwell time distributions from the indicated ND-BLM fusion pore conditions. The data from each replicated condition are pooled. **f** Opening and closing rates of ND-BLM fusion pores derived from the indicated closed and open dwell time analyses, respectively. Each condition was repeated at least five times on different days using fresh materials. Error bars represent standard error of the mean. Conditions were compared using the Student’s t-test. Within the bar graphs, ****, ***, **, * and ns represents p<0.0001, p<0.001, p<0.01, p<0.05 and a non-significant difference, respectively.

In line with Das *et al*. (2020), we validated that in the absence of Ca^2+^ (0.5 mM BAPTA), ND_3_-syt1 formed small and unstable fusion pores in all conditions (**Figs. 5b & 5c**). The addition of Ca^2+^ caused a significant increase in the current passing through ND_3_-syt1 fusion pores that form in the DOPS-containing BLM (**Figs. 5b & 5c**), indicative of an increase in pore size. Ca^2+^ also significantly increased the fraction of time that the ND_3_-syt1 pores remained in the open state in the DOPS-containing BLM (**Fig. 5d**). The peak of the open dwell time distribution was shifted to longer open times by >100-fold (**Fig. 5e**), compared to the BAPTA condition. Interestingly, we found that Ca^2+^ failed to trigger syt1-mediated fusion pore expansion in the DPPS-containing BLM (**Figs. 5b-5e**), presumably due to impaired membrane insertion. Specifically, the current passing through ND_3_-syt1 pores formed in the DPPS-containing BLM was unchanged by the addition of Ca^2+^; the open dwell time and the fraction of time that pores remained in the open state were also unaffected by Ca^2+^. Together, these data suggest that syt1 must penetrate the target membrane in order to drive stable fusion pore opening and dilation.

### Synaptotagmin 7 robustly penetrates ordered PS bilayers to dilate and stabilize fusion pores

To further correlate membrane penetration with fusion pore opening and dilation we aimed to purify and reconstitute full-length syt1 5W (FL-syt1 5W), and full-length syt7 (FL-syt7) into NDs to compare with the WT syt1 ND-BLM recordings, as the C2AB domains of syt1 5W and syt7 were able to efficiently bind (**Figs. 4c & 4f**) and penetrate (**Figs. 1, 2 and Supplementary Fig. 8**) into DPPS-containing bilayers. However, to date, the use of recombinant FL-syt7 has not been reported, presumably due to difficulty in purification. We were able to express and purify functional FL-syt7, using high volumes of starting material and a stringent protocol, which enabled us to examine the impact of FL-syt7 on fusion pores, as compared to FL-syt1 and FL-syt1 5W. We reasoned that if syt1 5W and syt7 C2AB domains were able to penetrate saturated PS bilayers, and if penetration is indeed critical for regulating pores, then both FL-syt1 5W and FL-syt7 should be capable of opening and dilating fusion pores that form in a DPPS-bearing BLM, in response to Ca^2+^. Indeed, increased hydrophobicity of the syt1 C2AB penetration loops was previously shown to enhance fusion pore expansion in HeLa cells expressing flipped t-SNAREs^56^ and in PC12 cells^57^. In line with enhanced penetration performance, compared to WT syt1, we found that FL-syt1 5W and FL-syt7 reconstituted into ND_3_ both yielded significant increases in fusion pore current in DPPS-bearing BLMs in the presence of Ca^2+^ (**Figs. 5b & 5c**). The addition of Ca^2+^ also significantly increased the fraction of time that ND_3_-syt1 5W and ND_3_-syt7 pores were in the open state (**Fig. 5d**) and the open dwell time distribution shifted to approximately 10-fold longer open times (**Fig. 5e**).

When examining the kinetics of fusion pore transitions in the presence of Ca^2+^, we found that the closing rates for ND_3_-syt1 fusion pores formed in the DPPS-BLM were significantly faster than for the DOPS condition (**Fig. 5f**, *upper panel*); FL-syt1 failed to stabilize the fusion pore open state when membrane penetration was impaired by using DPPS. Moreover, the closing rates of the ND_3_-syt1 fusion pores in the DPPS-containing BLM were also significantly faster than the penetration-competent ND_3_-syt1 5W and ND_3_-syt7 samples when Ca^2+^ was present (**Figs. 1, 5f and Supplementary Fig. 8**), further supporting the conclusion that stabilization of the open state is likely a direct consequence of C2-domain penetration into the target BLM. Interestingly, while Ca^2+^ enabled ND_3_-syt1 to significantly increase the fusion pore opening rate in the DOPS BLM, we found no difference in the opening rates between ND_3_-syt1, ND_3_-syt1 5W and ND_3_-syt7 in the DPPS condition (**Fig. 5f**, *lower panel* **and Supplementary Fig. 11**). The discrepancy in opening rates between the penetration competent samples, ND_3_-syt1 in DOPS, as well as ND_3_-syt1 5W and ND_3_-syt7 in DPPS, may suggest that C2-domain membrane penetration is preceded by SNARE zippering. Alternatively, the presence of DPPS at the fusion pore site may alter the conformation of the t-SNAREs^51^ or increase the local rigidity of the phospholipid bilayer^58^, to affect pore opening. Indeed, it was previously demonstrated that saturated acyl chains reduce SNARE-mediated fusion^59^. It is also noteworthy that in the ND-BLM BAPTA conditions, PC and PS (80% and 20%, respectively) are expected to be uniformly distributed throughout the BLM (**Supplementary Fig. 5a**). Therefore, SNARE-alone fusion pores could form in a PS-independent manner (i.e., by forming a malleable, DOPC rich fusion pore) and are thus less affected by the acyl chain structure of the PS constituent. However, in the DPPS BLM, Ca^2+^ is expected to cause the syts to act at the site of, or adjacent to, a rigid PS cluster (**Figs. 2 and Supplementary Fig. 5a**), which may affect the kinetics of pore opening. Regardless, upon pore opening, the penetration competent syts act to restrict closure and drive dilation.

The ND-BLM data presented thus far exclusively employed our newly developed method of forming planar lipid bilayers via drying and re-hydrating the lipids across the aperture of the BLM cup (**Fig. 5A**); this facilitated the incorporation of DPPS into the BLM and enabled us to study how syt penetration impacts fusion pore properties (**Fig. 5**). Next, we reverted to classical BLM painting methodology^41, 55^ to make a direct comparison of syt1 and syt7 when both proteins were capable of penetrating the target membrane in response to binding Ca^2+^. These experiments used ND_3_-syt1 and ND_3_-syt7 in association with DOPS-containing t-SNARE BLMs. Using this approach, syt7 continued to generate significantly larger fusion pores than syt1, but the kinetic analysis found no significant differences in open time fractions, or the closing and opening rates of the pores that are regulated by these two proteins (**Supplementary Fig. 12**). Thus, when membrane penetration is no longer a limiting factor, the functional differences in syt1 and syt7 are more subtle under these conditions.

## Discussion

During membrane fusion, discrete phospholipid bilayers are drawn together by proteins to overcome significant energy barriers and drive membrane merger. Recently, we described that an accessory protein, complexin, promotes SNARE-mediated membrane fusion by stabilizing curved intermediate structures that form during fusion via insertion of a C-terminal amphipathic helix at the fusion pore site^42^. Syt1 has also been suggested to promote fusion via membrane insertion^34, 37–39^. However, with the multitude of functions assigned to syt1, precisely how this molecule synchronizes Ca^2+^ influx into nerve terminals with the release of neurotransmitters remains unclear. In particular, the specific contribution of syt1 membrane penetration in triggering fusion has been difficult to disentangle from other interactions. For example, syt1 inhibits SV exocytosis in the absence of Ca^2+^ (a fusion clamp), presumably by binding SNAREs and preventing complete zippering^9^. In principle, Ca^2+^ binding could drive synchronized SV exocytosis simply by re-directing syt1 away from the *trans*-SNARE complex to release the fusion clamp and trigger fusion, potentially making membrane penetration inconsequential. Arguing against this idea, however, Ca^2+^•syt1 has been shown to directly drive SNARE complex assembly via PS binding^40^. Previous studies have examined the role of syt1 membrane penetration by mutagenesis to increase or decrease the hydrophobicity of the C2-domain penetration loops^39, 56, 60^. These studies support the importance of membrane penetration, but -as described above - these mutants may have unintended effects on protein function that confound conclusions. Namely, these mutations have been shown to affect the ability of syt1 to bind SNARE proteins^39^. Importantly, however, other mutagenesis studies de-coupled the effects of membrane penetration and SNARE-binding on exocytosis^14, 61^. It was demonstrated that altering the rigidity and/or rotational orientation between the tandem C2-domains of syt1 affected membrane penetration performance in correlation with exocytosis, without affecting SNARE-binding^14, 61^, but effects on interactions with other effectors could still not be ruled out.

To specifically assess the role of syt1 membrane penetration in the fusion reaction, we took an alternative approach by primarily working with WT proteins and then manipulating the composition of the target membrane to control syt1•membrane interactions. We found that syt1 cannot bind or penetrate phospholipid bilayers containing saturated PS (**Figs. 1-4**). We reasoned that this phenomenon could be exploited to isolate the role of membrane penetration from all other interactions to gain detailed insights into how syt1 triggers fusion. We also found that other C2-domain containing proteins (i.e. cPLA2, PKC and Doc2β) failed to penetrate bilayers containing saturated acyl chains (**Supplementary Fig. 3**). Surprisingly, however, we discovered that syt7 exhibited robust membrane penetration into bilayers containing either saturated or unsaturated PS (**Figs. 1 & 2**). This distinction enabled direct comparisons between syt1 and syt7, to tease apart the specific role of membrane penetration in syt function.

The distinct membrane penetration performance of syt1 and syt7, both experimentally (**Fig. 1**) and via MD simulations (**Fig. 2 and Supplementary Fig. 6**), then led us to investigate a mechanism that enables syt7 to act as a ‘super-penetrator’. We therefore examined the relative contributions of hydrophobic and electrostatic interactions in mediating syt•membrane binding. We demonstrate that Ca^2+^-dependent binding of C2-domains to phospholipid bilayers requires membrane penetration (**Fig. 4**); in isolation, electrostatic interactions are insufficient. Indeed, although Ca^2+^-bound syt1 binds to di-acylated anionic lipids with high affinity, syt1 does not associate with mono-acylated PS (lyso-PS)^62^, which originally suggested that syt1 membrane binding is a combination of electrostatic and hydrophobic interactions. Hydrophobic residues in the membrane penetration loops of both syt1 and syt7 are indeed critical for binding phospholipid bilayers (**Fig. 4**). Specifically, increasing syt1 hydrophobicity overcomes the defect in binding to saturated PS and slows the syt1•membrane disassembly kinetics (**Figs. 4e, 4f & 4i**). Conversely, reduced hydrophobicity renders syt7 susceptible to elevated salt (**Figs. 4g & 4h**) and speeds up the syt7-membrane disassembly kinetics (**Fig. 4i**). Curiously, the primary amino acid sequence of the syt1 and syt7 penetration loops (**Fig. 4d**) failed to predict the binding properties of these C2-domains, suggesting that membrane binding is more complex than simply a sum of their charged and hydrophobic residues.

After finding that syt1 membrane binding can be controlled by manipulating the PS acyl chain structure, and that syt7 and a gain-of-function syt1 (syt1 5W) can overcome the saturated PS binding defect observed with WT syt1, we proceeded to test these three variants functionally, using the ND-BLM approach (**Supplementary Fig. 10**). To our knowledge, this is the first reported successful purification, reconstitution and functional testing of FL-syt7. By comparing syt action upon membranes containing DOPS or DPPS, we aimed to determine whether membrane penetration is required for syt1/7 to drive fusion pore opening and dilation. To address this, a new BLM method was developed in order to enable the formation of planar lipid bilayers containing all the lipids used in this study (**Fig. 5a**). This method of incorporating DPPS in the BLM was critical as the distinct penetration performances of syt1 and syt7 are otherwise subtle (**Supplementary Fig. 12**). As predicted, we observed that stable fusion pore opening, and dilation, were directly correlated with the ability of syts to penetrate membranes. Specifically, when C2-domains are incapable of penetrating a bilayer, fusion pore stabilization and dilation is abrogated. Conversely, conditions that permit the syts to penetrate membranes in response to Ca^2+^ (syt1 and syt7 into DOPS, or syt1 5W and syt7 into DPPS) result in larger, more stable fusion pores (**Fig. 5 and Supplementary Fig. 12**). Hence, syt membrane penetration activity directly mediates, at least in part, the regulation of fusion pores.

Kinetic analysis revealed that membrane penetration competent syts significantly reduced fusion pore closing rates, thereby keeping pores in the open state for longer periods (**Fig. 5f**). Notably, the ND-BLM system traps fusion pores in a reversible intermediate state, due to the rigid scaffold that surrounds the NDs. However, *in vivo*, no such restricting scaffold is present; the penetration action of the syts in neurons may instead favor a one-way reaction by stabilizing the nascent fusion pore, reducing the propensity to reverse towards closure, and supporting pore dilation. In this view, Ca^2+^ binding by syts directs the C2-domains to penetrate the target membrane^63, 64^, thus lowering the energy barrier for full fusion of SV with the plasma membrane.

In chromaffin cells syt1 and syt7 both localize to the surface of secretory granules where they were reported to differentially activate exocytosis^65^. These two syt isoforms were shown to partially segregate into non-overlapping granule pools that exhibit distinct modes of exocytosis. Specifically, membrane depolarization triggered rapid fusion and full collapse of syt1-granules with the plasma membrane, while syt7-granules released their encapsulated contents slowly and appeared to restrict full vesicular collapse^65^. Under our current experimental conditions, the ND-BLM results failed to detect these distinct phenomena. This may be attributed our use of small (13 nm) NDs that have limited dilation capacity. Future studies that aim to specifically examine the fusion pore dilation step may instead exploit larger NDs to reveal unique effects of syt1 and syt7 on dilation.

It is noteworthy that, while syt7 was determined herein to exhibit more robust binding and penetration into lipid bilayers, as compared to syt1, recent findings suggest that syt7 might not act solely as a Ca^2+^ sensor for exocytosis in all cell types^21, 23, 32^. For example, in neurons, syt7 functions as a dynamic, Ca^2+^-regulated SV docking protein on the axonal plasma membrane that feeds docked vesicles to Doc2α, another slow sensor that triggers asynchronous neurotransmitter release^27^. In this context, with highly curved ∼42 nm SVs, the binding and penetration of syt7 into the vesicle bilayer might mediate its recently described activity-dependent docking function. We note that, syt7 has also been reported to be a direct exocytic Ca^2+^ sensor residing on lysosomes, contributing to lysosome•plasma membrane fusion^28^ Yet another study suggested that syt7 on dense core vesicles plays a role in docking or priming, rather than fusion per se^32^. Clearly, additional work is needed to clarify the function of syt7, but an appealing idea is that syt7 functions differently when targeted to the presynaptic membrane in neurons versus when it is targeted to dense core vesicles or lysosomes. We also note that roles for syt7 in dense core vesicle docking/priming and fusion are not mutually exclusive.

In this study, we reconstituted FL-syt7 into NDs to facilitate a direct comparison with syt1 using the ND-BLM approach. Our main goals were to assess whether syt7 can directly modulate fusion pores, and if so, how membrane penetration influences this regulation. We established that syt7 exhibits unusually robust membrane binding properties and that membrane penetration is indeed a critical step that enables both syt1 and syt7 to directly regulate fusion pores in response to binding Ca^2+^. We note that interrogation of syt1 and syt7 function in the ND-BLM experiment mimics, in a sense, dense-core granule exocytosis, due to both isoforms residing in the ND. Ongoing studies will incorporate FL-syt7 into the BLM and syt1 in the ND, to model SV biology, *in vitro,* with increased complexity. Moreover, with recombinant FL-syt7 readily in hand, future studies will use FL-syt7 to reconstitute the SV docking step, with the goal of developing an *in vitro* model to build a more complete understanding of the SV cycle.

## Author contributions

K.C.C., Q.C. and E.R.C. designed the study; K.C.C., T.M., N.M., L.W., Y.L. and D.D performed experiments and analyzed data; K.C.C. and E.R.C. wrote the manuscript; E.R.C. and Q.C provided funding.

## Acknowledgements

We are grateful to the other members of the Chapman lab and M.B. Jackson for providing critical feedback on this project. We also especially thank C. Greer for editorial contributions and D. Larson for management of cDNA records. This work was supported by National Institutes of Health grants MH061876 and NS097362 (E.R.C.) and the National Science Foundation grant NSF-DMS1661900 (Q.C.) Computational resources from the Extreme Science and Engineering Discovery Environment (XSEDE(49)), which is supported by NSF grant number ACI-1548562, are greatly appreciated; part of the computational work was performed on the Shared Computing Cluster which is administered by Boston University’s Research Computing Services (URL: www.bu.edu/tech/support/research/). E.R.C. is an Investigator of the Howard Hughes Medical Institute (HHMI). This article is subject to HHMI’s Open Access to Publications policy. HHMI lab heads have previously granted a nonexclusive CC BY 4.0 license to the public and a sublicensable license to HHMI in their research articles. Pursuant to those licenses, the author-accepted manuscript of this article can be made freely available under a CC BY 4.0 license immediately upon publication.

---

**Supplementary Fig. 1.**
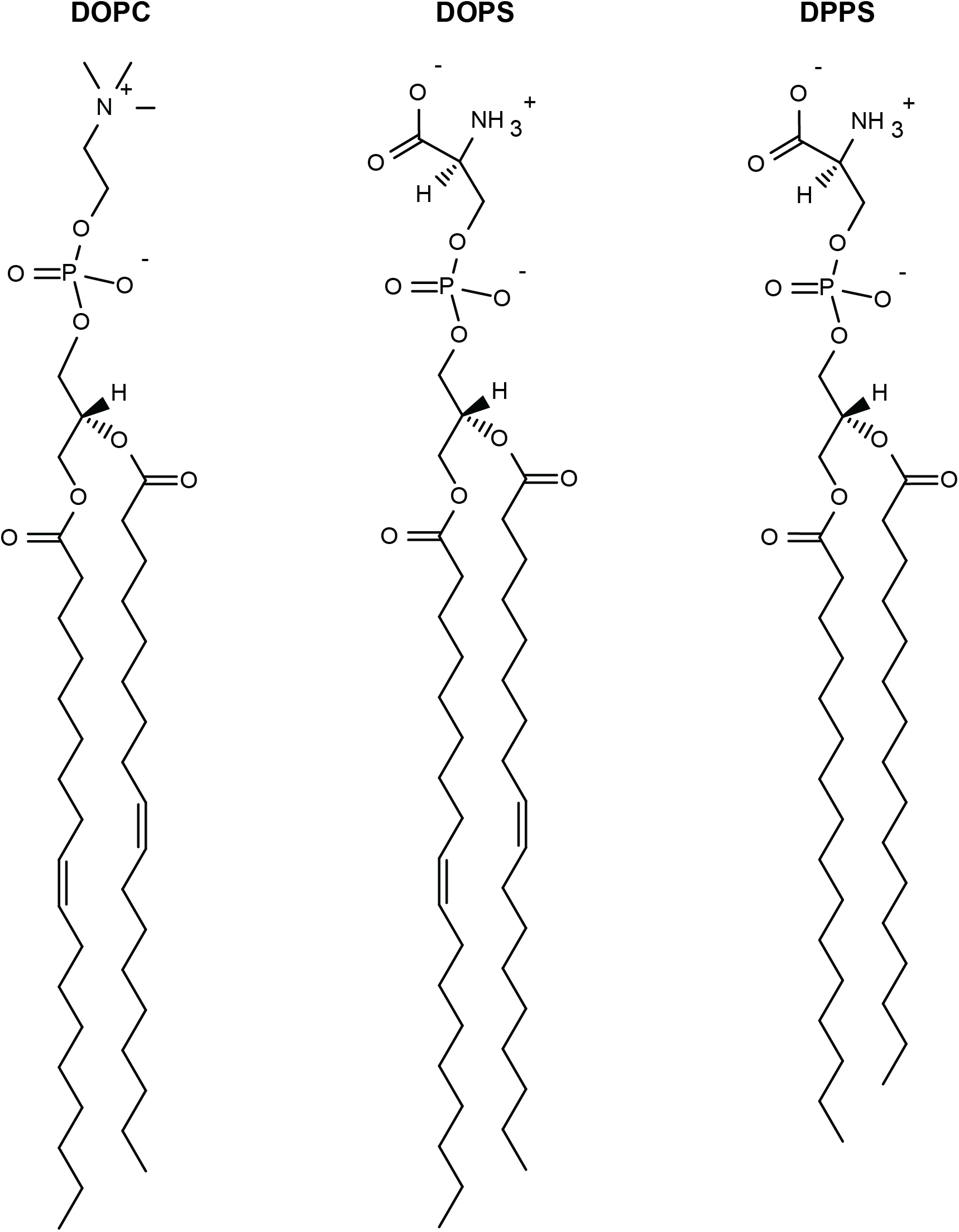
Phospholipid structures. Chemical structures of the phospholipids used in this study

**Supplementary Fig. 2.**
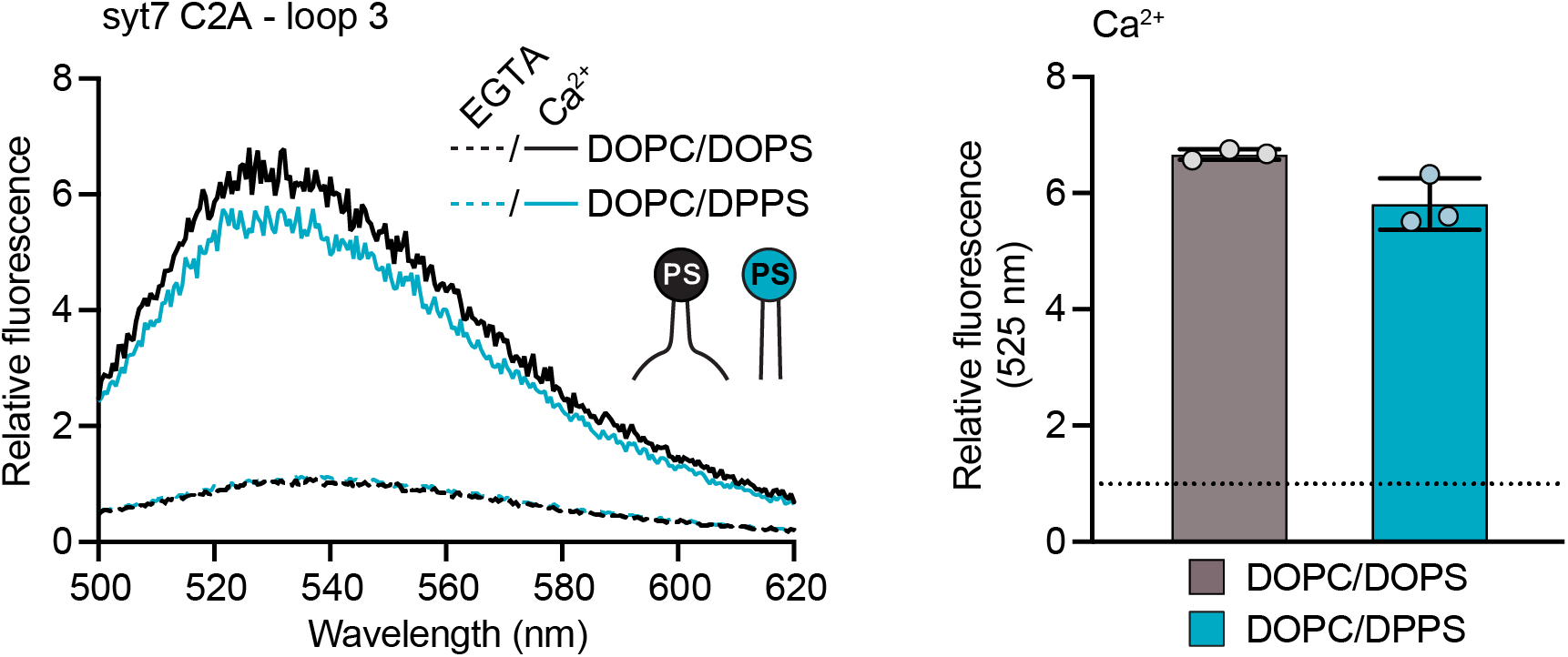
The C2A domain of syt7 C2AB equally penetrates bilayers containing DOPS and DPPS. Representative fluorescence emission spectra (*left panel*) of NBD labelled syt7, in the presence (solid lines) and absence (dotted lines) of Ca^2+^, and liposomes composed of 80:20 DOPC/DOPS (black) or DOPC/DPPS (blue). The syt7 C2AB domain is labeled on loop 3 of the C2A domain at position 229. Quantification of NBD-syt7 C2AB fluorescence emission at 525 nm in the presence of Ca^2+^ and liposomes composed of DOPC/DOPS (black) or DOPC/DPPS (blue), *right panel*. The data are normalized to the EGTA condition, shown as a horizontal black dotted line. Each condition was repeated three times on different days using fresh materials. Error bars represent standard error of the mean.

**Supplementary Fig. 3.**
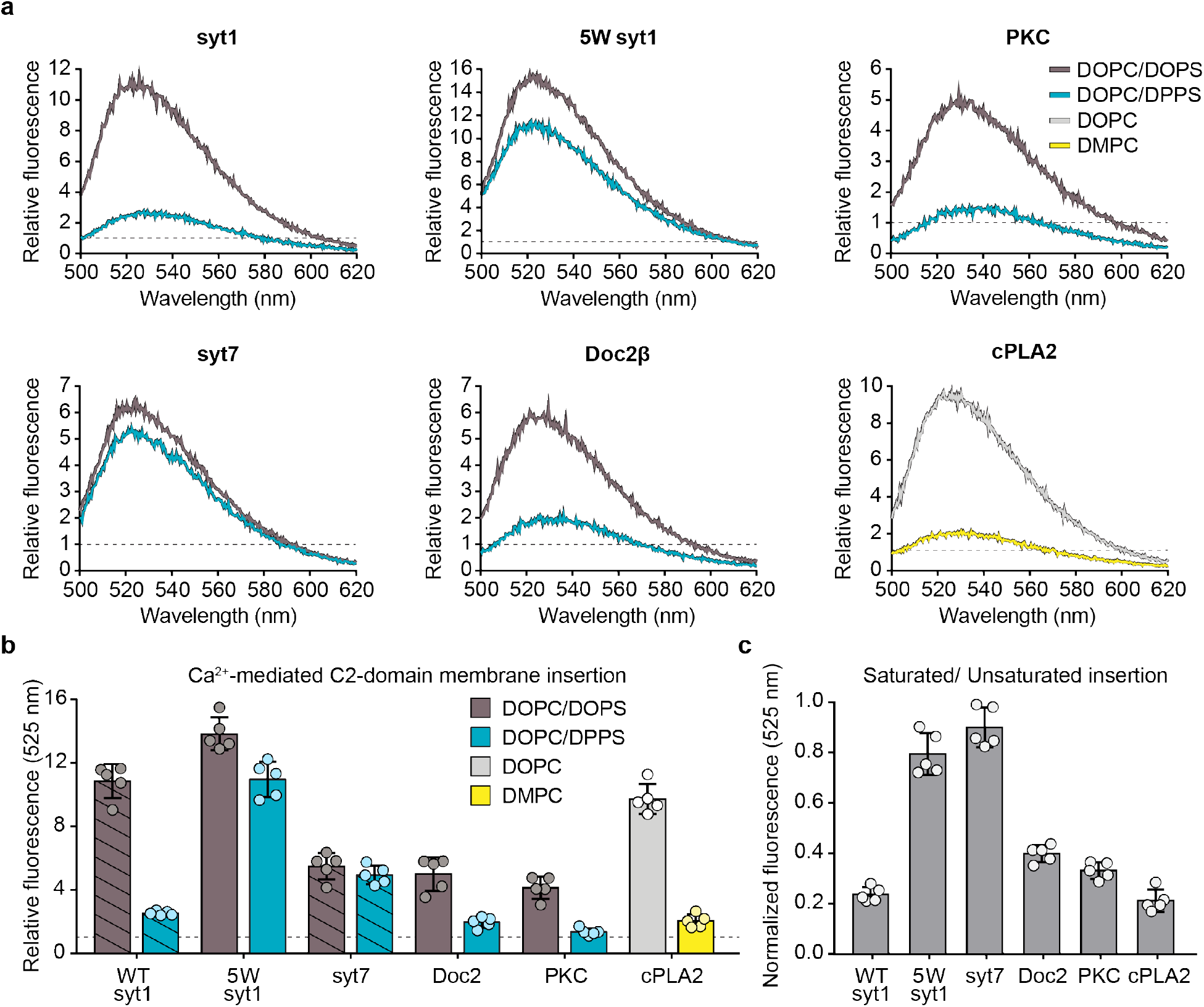
Doc2β, PKC and cPLA2 fail to efficiently penetrate bilayers with saturated acyl chains. **a** Representative fluorescence spectra from NBD labeled Doc2β, PKC and cPLA2 in the presence and absence of Ca^2+^. The NBD fluorescence in EGTA is represented as a black dotted line. The Doc2β and PKC experiments were performed with liposomes composed of DOPC/DOPS or DOPC/DPPS (80:20); the cPLA2 experiments were performed with liposomes composed of DOPC or DMPC, due to the inherent lack of PS binding specificity. The syt1 and syt7 C2AB traces are reproduced from Fig. 1 for comparison. **b** Quantification of replicated experiments described in *panel a.* Replicates were performed at least five times on separate days with fresh lipid samples. The fluorescence intensity at 525 nm was extracted from each spectrum, normalized to the EGTA condition and plotted. The horizontal dotted line represents the normalized fluorescence intensity the EGTA conditions. The syt1 and syt7 data are reproduced from Fig. 1 for comparison and emphasized with diagonal lines. **c** Normalized NBD fluorescent intensities at 525 nm from data represented in *panel b*. The fluorescence in the saturated acyl chain conditions were divided by the condition containing all unsaturated acyl chains. Error bars represent standard error of the mean.

**Supplementary Fig. 4.**
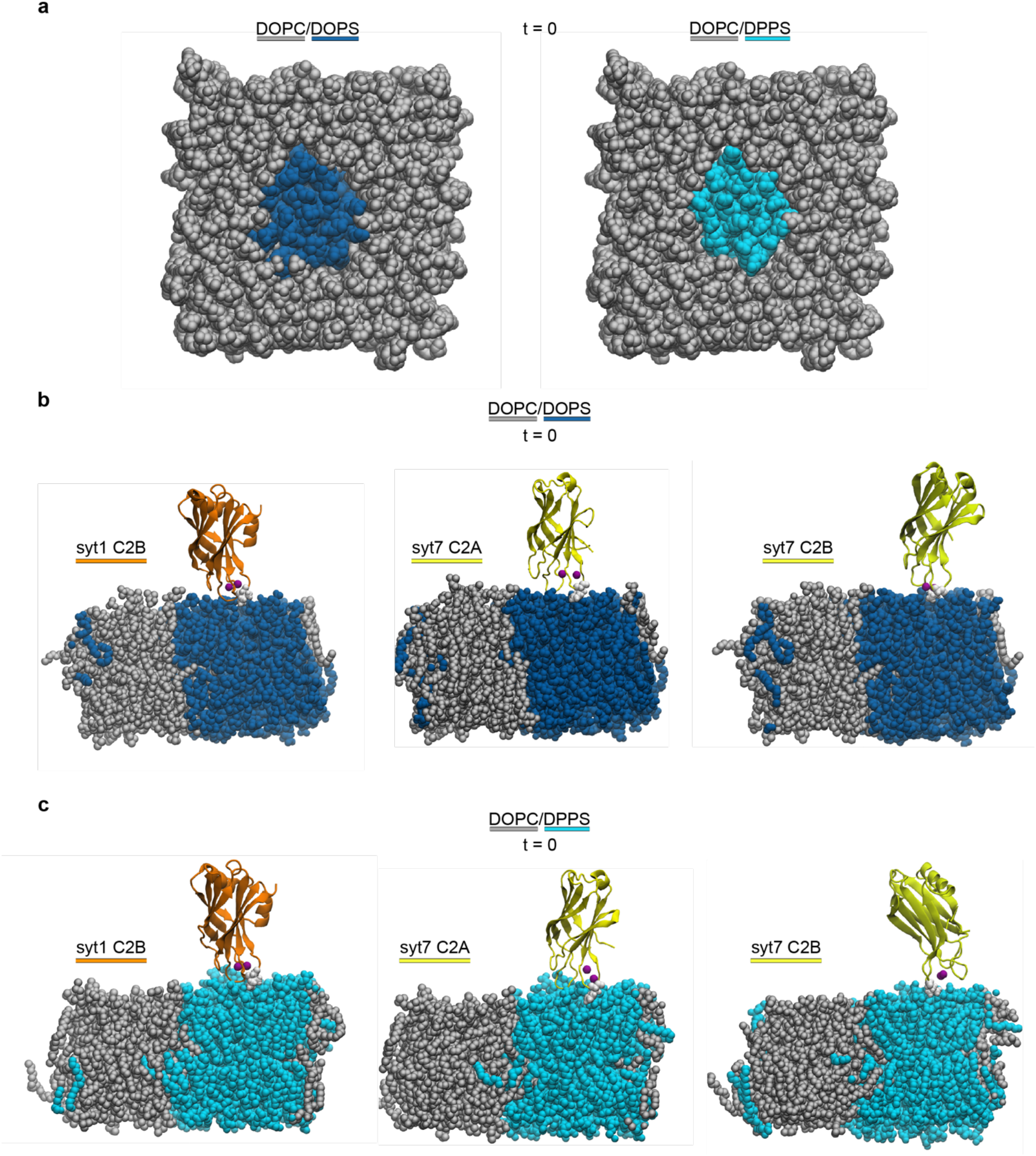
Initial configurations of MD simulations. **a** Time zero snapshots of MD simulations that examine the clustering behavior of DOPS (*left*) and DPPS (*right*) among DOPC lipids (grey). **b** Time zero snapshots of MD simulations comparing syt1 C2B (shown in orange) domain with syt7 C2A and C2B domains (yellow) positioned on bilayers containing 20% DOPS. **c** Time zero snapshots of MD simulations comparing syt1 C2B (shown in orange) domain with syt7 C2A and C2B domains (yellow) positioned on bilayers containing 20% DPPS.. Corresponding end-point snapshots for syt1 and syt7 C2B domains are found in Figure 2. The syt7 C2A end-point snapshots are shown in Supplementary Fig. 6.

**Supplementary Fig. 5.**
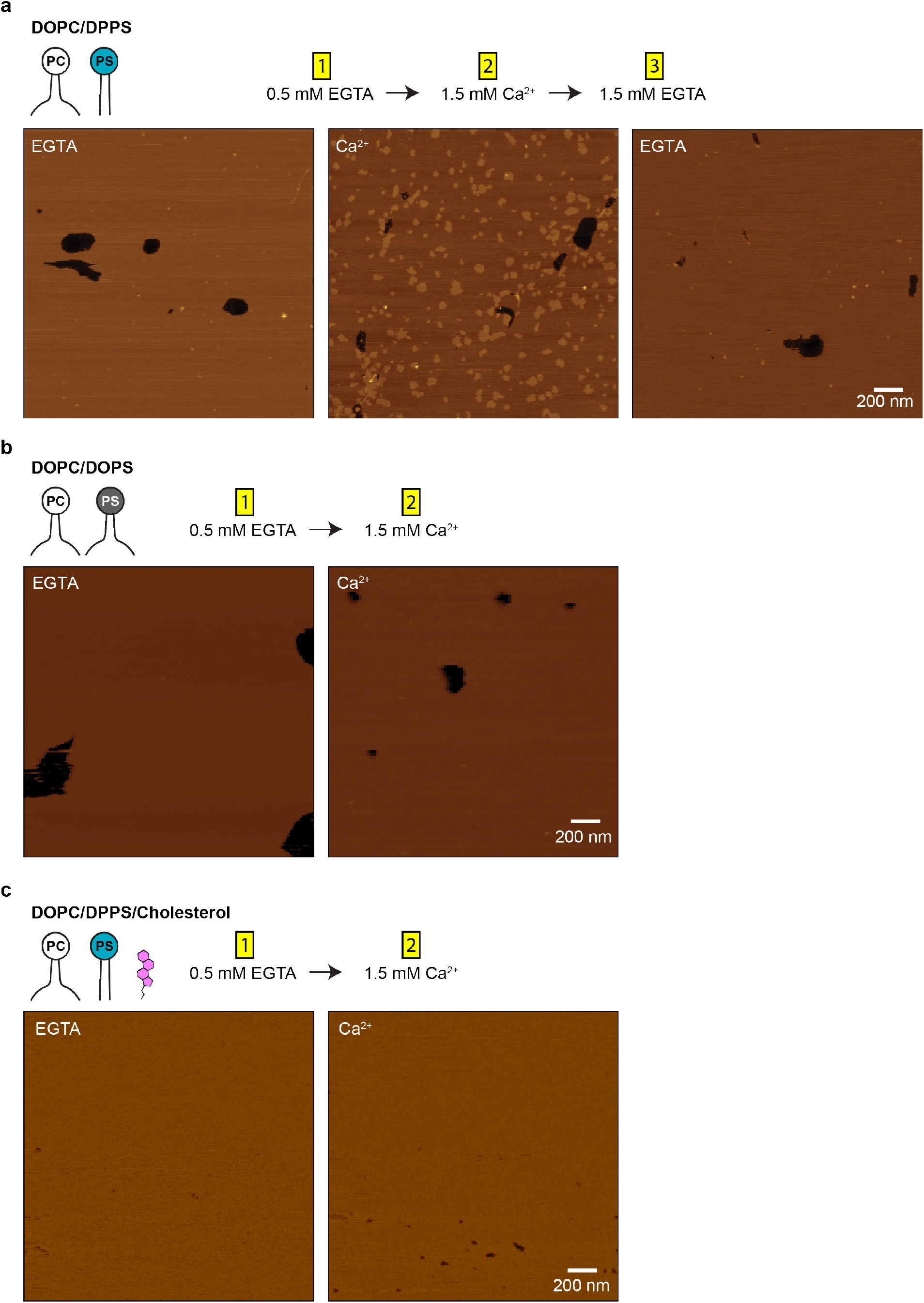
Ca^2+^ causes DPPS clustering in lipid bilayers. **a** AFM imaging of supported lipid bilayers on mica composed of DOPC/DPPS (80:20) with and without Ca^2+^. **b** AFM imaging of supported lipid bilayers on mica composed of DOPC/DOPS (80:20) with and without Ca^2+^. **c** AFM imaging of supported lipid bilayers on mica composed of DOPC/DPPS/cholesterol (56:14:30) with and without Ca^2+^.

**Supplementary Fig. 6.**
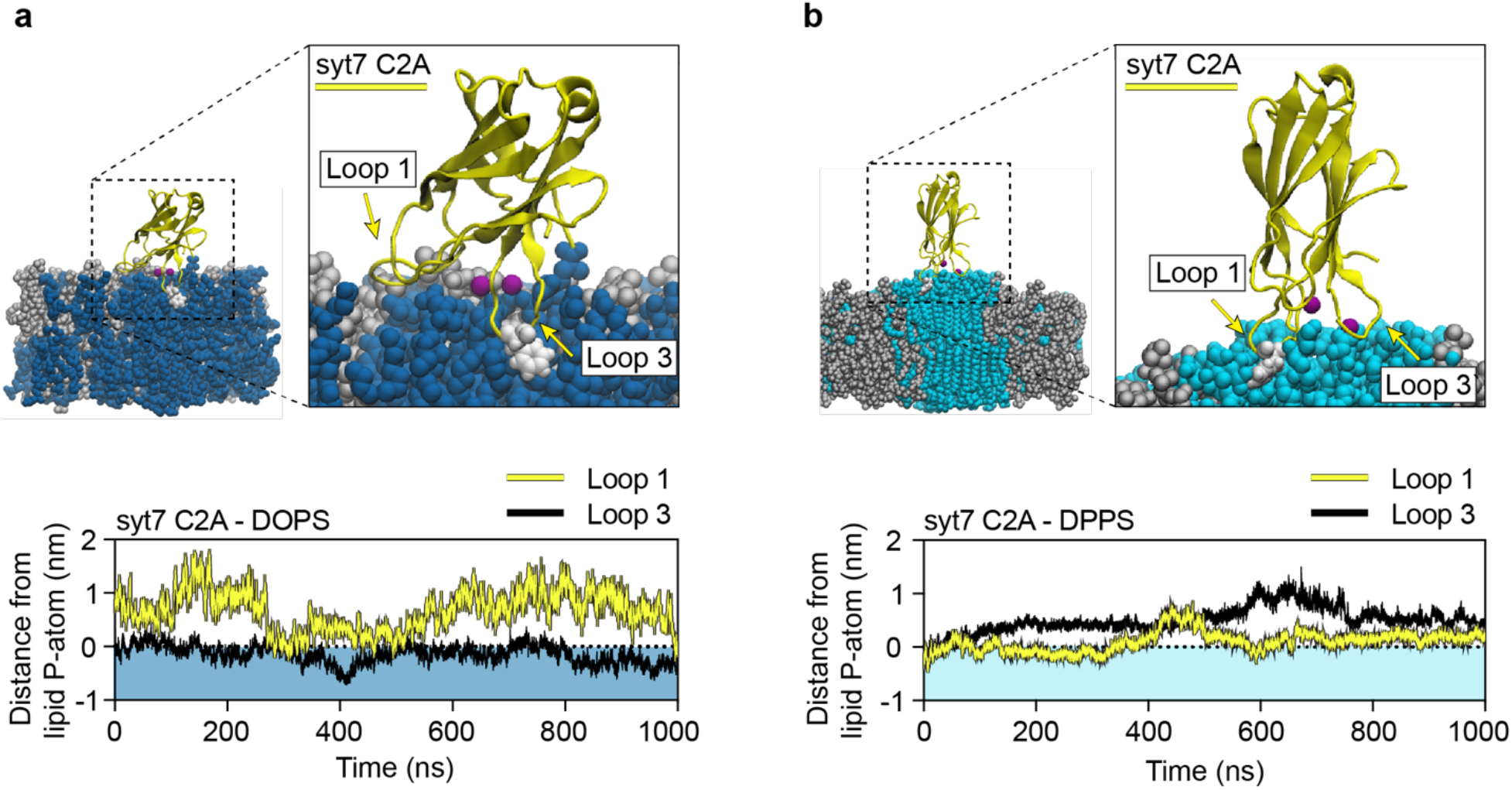
Molecular dynamics simulations of syt7 C2A•membrane interactions. **a** End point (1000 ns) MD simulations snapshot showing the syt7 (yellow) C2A domain interacting with a lipid bilayer composed of DOPC/DOPS. The loop residue that achieved the deepest depth of penetration, F229, is emphasized in white. **b** End point (1000 ns) MD simulations snapshot showing the syt7 (yellow) C2A domain interacting with a lipid bilayer composed of DOPC/DPPS. The loop residue that achieved the deepest depth of penetration, F167, is emphasized in white. Quantification of loop 1 and loop 3 depth from syt7 C2A into the DOPS-containing (blue shading) and DPPS-containing (cyan shading) bilayers are shown in the lower panels. In each case, the loop depth is normalized relative to the position of the lipid phosphate group, indicated by a horizontal dotted line.

**Supplementary Fig. 7.**
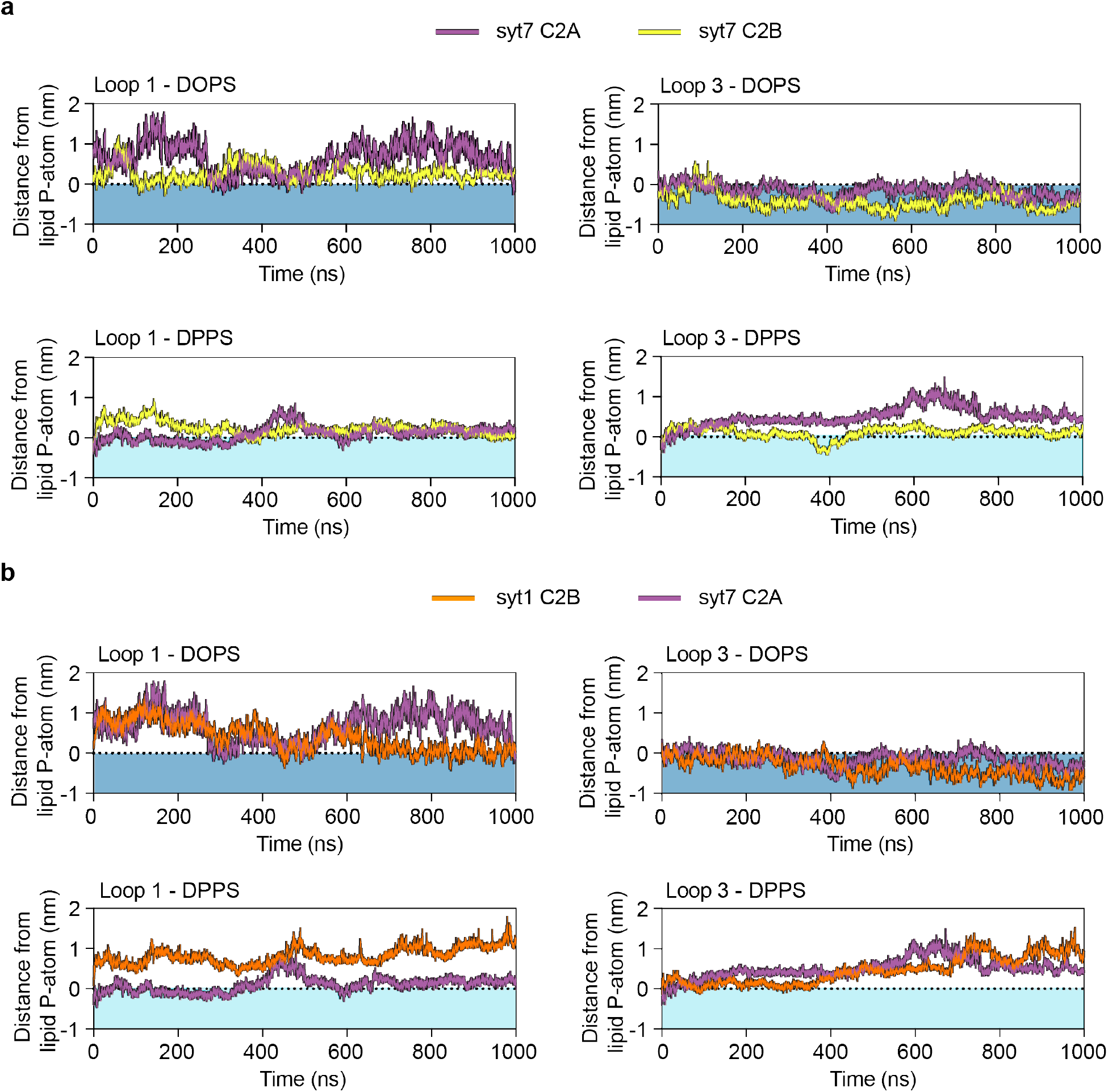
MD simulations comparison of membrane penetration depth between syt7 C2A and syt1 or syt7 C2B domains. **a** Quantification of loop 1 and loop 3 depth of syt7 C2A (purple) and syt7 C2B (yellow) domains into the DOPS-containing (blue shading) and DPPS-containing (cyan shading) bilayers. **b** Quantification of loop 1 and loop 3 depth of syt1 C2B (orange) and syt7 C2A (purple) domains into the DOPS-containing (blue shading) and DPPS-containing (cyan shading) bilayers. These data are re-plotted quantifications from Fig. 2 and Supplementary Fig. 6.

**Supplementary Fig. 8.**
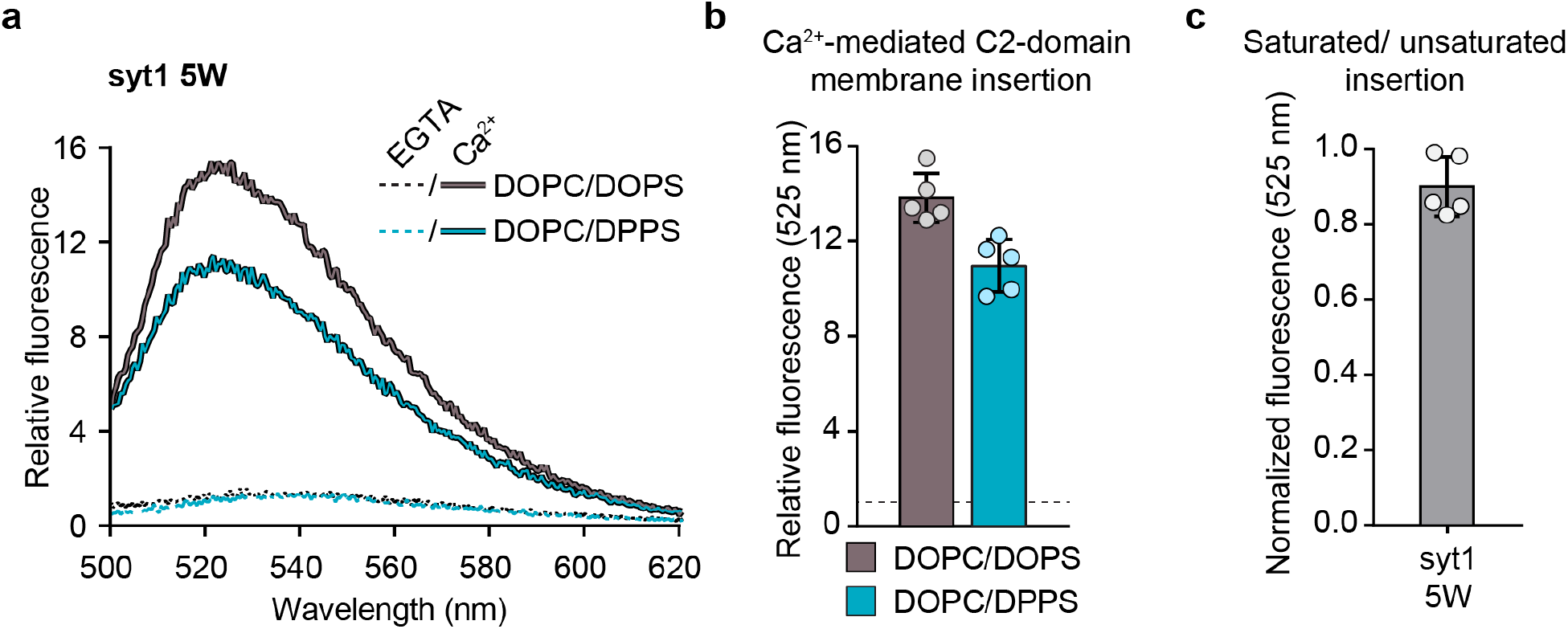
Tryptophan substitutions in the penetration loops enables syt1 to penetrate bilayers that harbor DPPS. **a** Representative fluorescence spectra of NBD-labelled syt1 5W C2AB in 0.2 mM EGTA (dotted lines) or 0.5 mM free Ca^2+^ (solid lines) and with liposomes composed of DOPC/DOPS or DOPC/DPPS (80:20). **b** Quantification of replicated experiments depicted in *panel a*. The normalized fluorescence intensity at 525 nm in Ca^2+^ is shown relative to the EGTA condition. **c** Normalized NBD fluorescent intensity at 525 nm from data represented in *panel b*. The fluorescence in the saturated acyl chain conditions were divided by the condition containing all unsaturated acyl chains. Each condition was repeated five times on different days using fresh materials. Error bars represent standard error of the mean.

**Supplementary Fig. 9.**
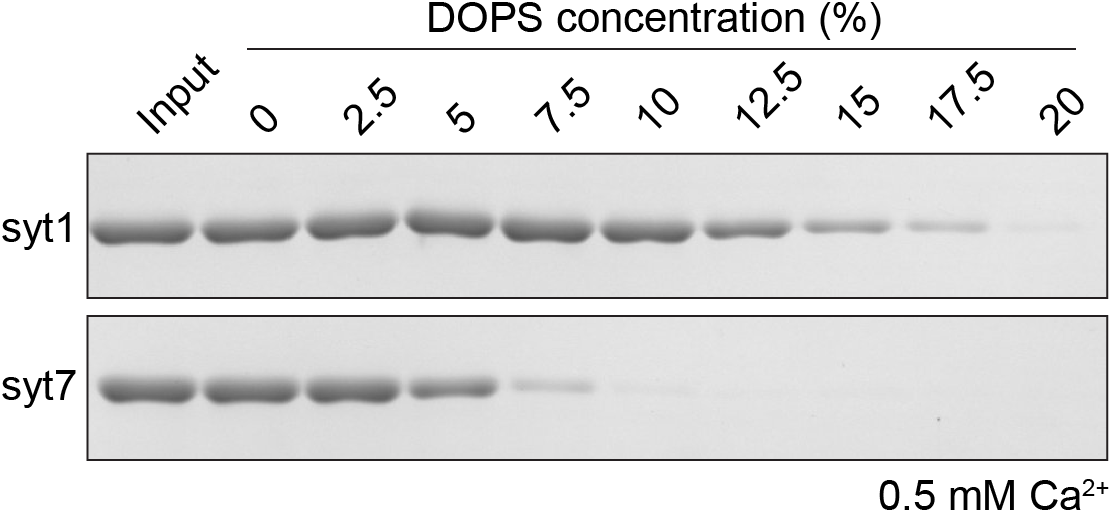
Syt7 displays higher affinity for PS-bearing liposomes than syt1. Representative Coomassie stained SDS-PAGE gel of a co-sedimentation assay (described in Fig. 4a) with syt1 and syt7 C2AB domains in the presence of Ca^2+^ and liposomes with increasing mole fractions of PS. Depletion of protein from the samples indicates binding to liposomes. Input refers to the protein-only control sample.

**Supplementary Fig. 10.**
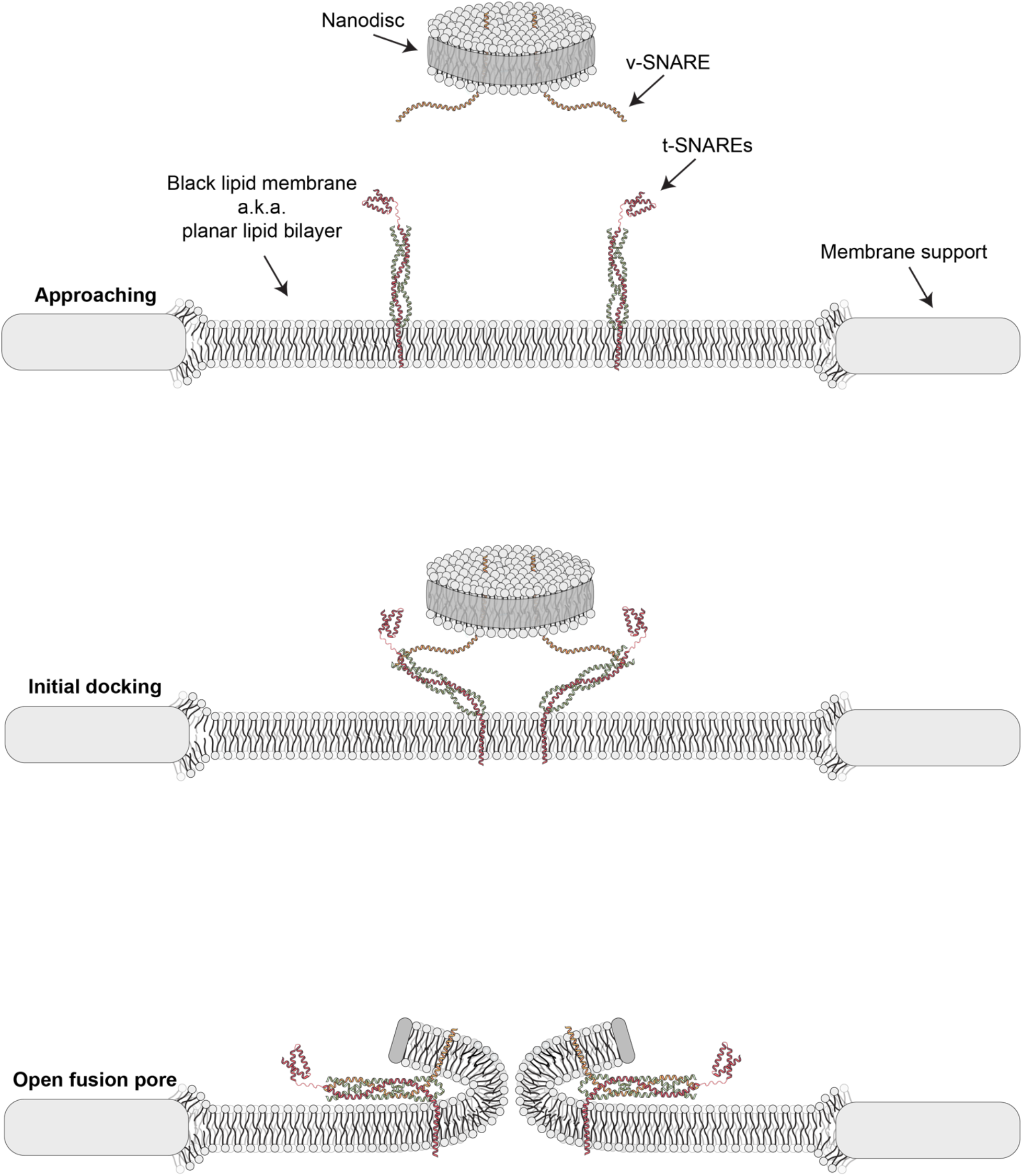
Assembly steps for fusion pore formation during nanodisc-black lipid membrane electrophysiology experiments. An illustration showing the sequence of events leading up to the formation of a fusion pore formed between a v-SNARE nanodisc and a t-SNARE black lipid membrane.

**Supplementary Fig. 11.**
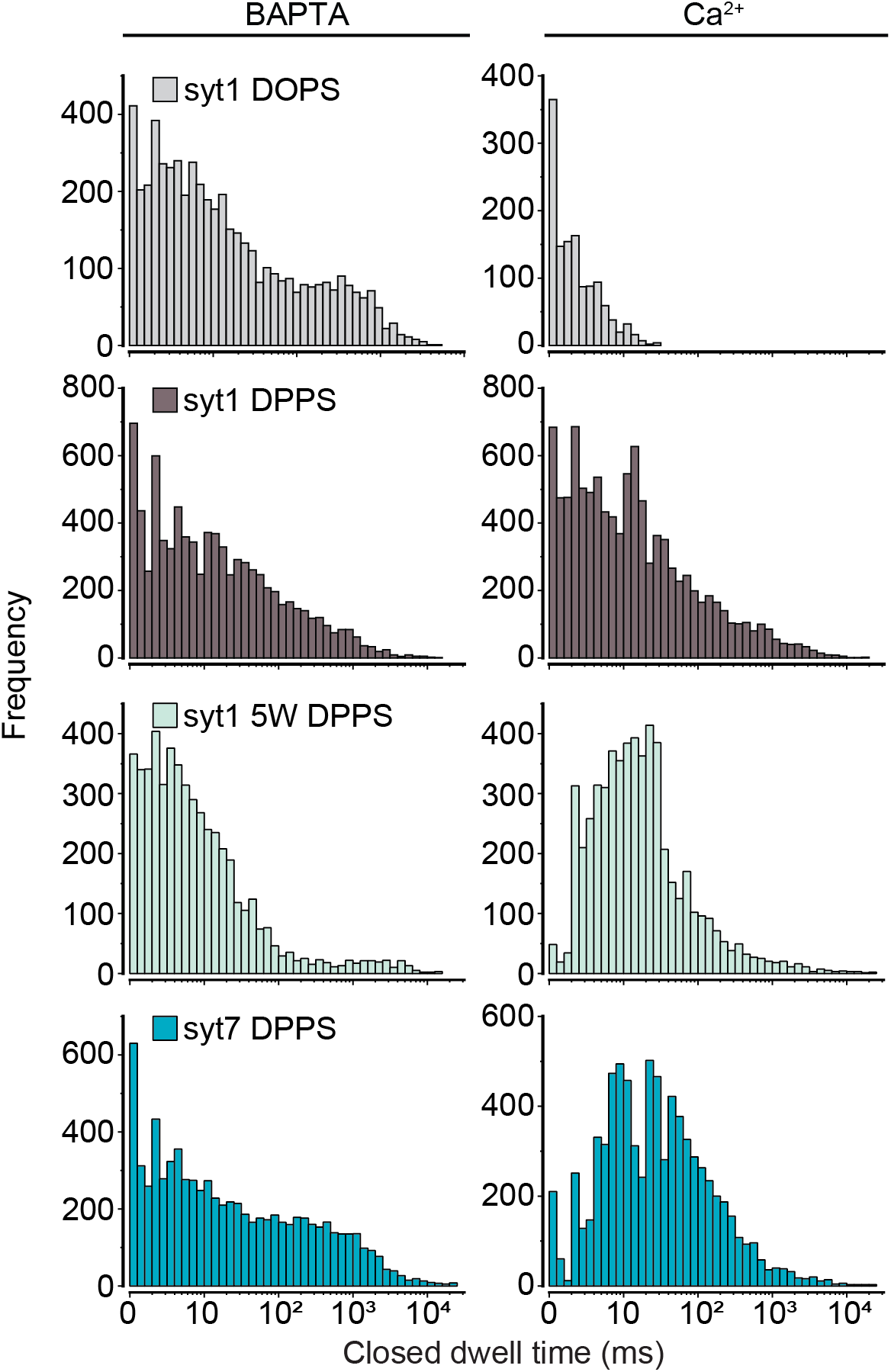
In the presence of Ca^2+^ and ND_3_-syt1 5W or ND_3_-syt7, SNARE-mediated fusion pores exhibit long closed dwell times when the BLM harbors DOPC/DPPS. Closed dwell time distributions generated and pooled from each of the indicated ND-BLM conditions, in 0.5 mM BAPTA or 0.5 mM free Ca^2+^. Associated data are found in Fig. 5.

**Supplementary Fig. 12.**
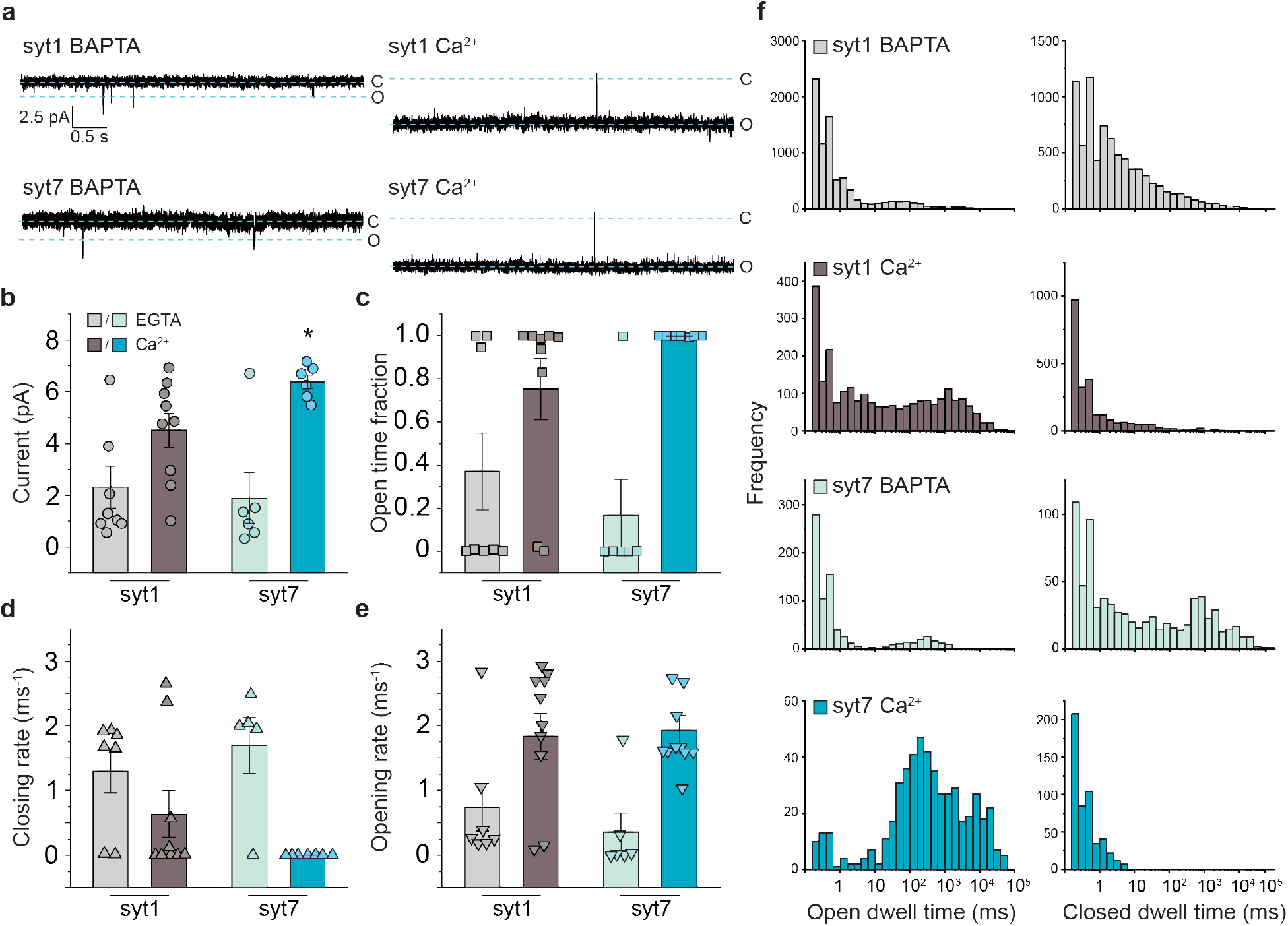
The differential effects of syt1 and syt7 on the properties of fusion pores are subtle in membranes containing DOPS. Results of ND-BLM experiments performed using classical BLM methodology with membranes containing 20% DOPS. **a** Representative ND-BLM traces from fusion pores formed by ND_3_-syt1 and ND_3_-syt7 in 0.5 mM BAPTA or 0.5 mM free Ca^2+^. **b** Quantification of the current passing through ND-BLM fusion pores under the indicated conditions. **c** Quantification of the fraction of time that ND-BLM fusion pores remained in the open state under the indicated conditions. **d** Closing rates of ND-BLM fusion pores derived from the indicated open dwell time analyses. **e** Opening rates of ND-BLM fusion pores derived from the indicated closed dwell time analyses. **f** Open and closed dwell time distributions from the indicated ND-BLM fusion pore conditions. The data from each replicated condition are pooled. Each condition was repeated six times on different days using fresh materials. Error bars represent standard error of the mean. The asterisk (*) in *panel b* indicates a significant difference between syt1 and syt7 in the 0.5 mM free Ca^2+^ condition was determined via Student’s t-test. All other comparisons between syt1 and syt7 throughout the figure are not significantly different.

Supplementary movie 1. **MD simulation of DOPS clustering within a DOPC bilayer.** A representative MD simulation showing a cluster of DOPS (blue) lipids within a DOPC bilayer (grey). The DOPS was placed at the center of the bilayer and allowed to equilibrate with the DOPC for 450 ns.

Supplementary movie 2. **MD simulation of DPPS clustering within a DOPC bilayer.** A representative MD simulation showing a cluster of DPPS (cyan) lipids within a DOPC bilayer (grey). The DPPS was placed at the center of the bilayer and allowed to equilibrate with the DOPC for 450 ns.

## Methods

### Protein purification

Recombinant proteins were produced in *E. coli* (BL21) and purified using Ni-NTA or TALON metal affinity agarose resin as previously described^42^. The cDNAs for FL-syt1, syt1 C2AB (residues 96-421; I367C), FL-syt1 5W (M173W, F231W, F234W V304W, Y364W, I367C), syt1 5W C2AB, cPLA2 (Y96C), PKC (L249C), FL-syt7, syt7 (L361C) and Doc2β (G361C) were cloned into a pET-SUMO vector and purified as SUMO fusion proteins. pET-SUMO-hGSDMD was a gift from Hongbo Luo (Addgene plasmid # 111559 ; http://n2t.net/addgene:111559 ; RRID:Addgene_111559); syt1 and PKC were provided by Tom Sudhof; syt7 was provided by Mitsunori Fukuda; Doc2β was provided by Matthijs Verhage; cPLA2 was provided by Roger Williams. With the exception of cPLA2 and PKC, the SUMO domain was cleaved from the protein of interest by incubation with 0.5 μM SENP2 protease overnight at 4 ⁰C. cPLA2, PKC, syb2, MSP1E3D1 and the SNAP-25B/syntaxin1a heterodimer were eluted from the Ni-NTA or TALON resin by 500 mM imidazole, followed by running the sample through a PD-10 desalting column to remove the imidazole. pMSP1E3D1 was a gift from Stephen Sligar (Addgene plasmid # 20066 ; http://n2t.net/addgene:20066 ; RRID:Addgene_20066); syb2 was provided by James Rothman; SNAP25B was provided by M.C Wilson; syntaxin1a was provided by Richard Sheller. SNAP25B and syntaxin1a were previously cloned together into the pRSFDuet-1 (Novagen)^66^. Proteins that contained a transmembrane domain were purified with the addition of 0.9% CHAPS in the buffers. The protein concentrations were determined by Coomassie stained SDS-PAGE and compared with a range of BSA standards.

### NBD labeling

The cPLA2 and PKC C2 domains were engineered to harbor a single cysteine residue in loop 3 of their single C2 domains, while syt1, syt1 5W, syt7 and Doc2β C2AB domains had a single cysteine in loop 3 of their C2B domains. The mutations for the single cysteine variants are listed above in the protein purification section. To conjugate NBD to the cysteine thiol group, 50 μM of each protein was incubated with at least 10-fold excess iodoacetamide-NBD overnight at 4 ℃, followed by removal of free dye using a PD-10 desalting column. Protein labeling efficiency (>80 %) was calculated using the Beers-Lambert law (A = εcl) with the NBD extinction coefficient of 25000 M^-1^ cm^-1^ at 480 nm to determine NBD content, and this value was compared to the protein concentration.

### Preparation of unilamellar vesicles

Large unilamellar vesicles (LUVs) were prepared by pipetting chloroform suspended lipids into a glass tube, followed by drying the sample under a stream of nitrogen gas. After formation of a thin lipid film, the sample was then placed in a lyophilizer for at least 60 minutes to remove residual organic solvent. The lipids were hydrated with 25 mM HEPES, 100 mM KCl pH 7.4 and placed at 50 ⁰C to aid in resuspension, followed by vortexing to generate multilamellar vesicles. The sample was then repeatedly passed through a 100 nm filter using a mini extruder system (Avanti Polar Lipids) to result in uniform LUVs.

### Co-sedimentation assay

One hundred nm LUVs (0.5 mM) composed of DOPC/DOPS or DOPC/DPPS (80:20) were mixed with 2.5 µM of the indicated C2AB proteins in 0.2 mM EGTA or 0.5 mM free Ca^2+^. The LUVs, or protein-LUV complexes, were then pelleted by ultracentrifugation at 160,000 x g in a TLA-100 rotor (Beckman). The unbound fraction of C2AB in the supernatants was quantified by densitometry analysis of Coomassie blue stained SDS-PAGE gels. For experiments with increasing salt, the samples received progressive increases in NaCl.

### Atomic force microscopy

Supported lipid bilayers composed of DOPC/DOPS, DOPC/DPPS (80:20) or DOPC/DPPS/Cholesterol (56:14:30) were prepared by first generating 1 mM LUVs, as described above, followed by incubating a 10-fold diluted sample with freshly cleaved mica discs in a liquid cell. The AFM imaging of the supported lipid bilayers was performed using an Agilent 5500 Scanning Probe Microscope, as previously described^67^.

### Molecular dynamics simulations

The syt1 C2B, syt7 C2A and syt7 C2B protein crystal structures were taken from the PDB database (PDB ID 1K5W, 2D8K and 3N5A, respectively). The lipid models were generated using CHARMM-GUI software^68^. The CHARMM36 forcefield^69^ was used to model all the components of the system. To check the stability of PS clusters, we first built a cylindrical cluster containing 32 DPPS or DOPS lipids. This PS cluster was then inserted into the center of a DOPC bilayer (**Supplementary Fig. 4a**). The lipid bilayer was then solvated using TIP3P water box of (∼9.4 × 9.4 × 8.0 nm^3^) and 32 Ca^2+^ and 32 Cl^-^ ions were added to the system. The solvated system was first energy-minimized using conjugate gradient method to remove any bad contacts between the solvent and solute atoms. This step was followed by a short NVT simulation in which phosphorous atoms of the lipid heads were restrained. The NVT equilibrated system was then subject to restraint-free NPT equilibration at the atmospheric pressure and room temperature. The temperature and pressure of the system were controlled using Nose-Hoover thermostat with a time constant of 1 ps and Parrinello-Rahman barostat with a time constant of 5 ps, respectively. The covalent bonds involving light hydrogen atoms were constrained using LINCS algorithm to enable the simulation time step of 2 fs. Periodic boundary conditions were enforced in all three directions and the long-range electrostatic interactions were calculated using particle-mesh-Ewald method. The short-ranged van der Waals forces was smoothly decayed to zero in the range of 1.0 to 1.2 nm using a switching function. All simulations were carried out using the Gromacs suite of programs^70^.

### Stopped-flow rapid mixing

Rapid mixing was performed using a SX-18.MV stopped-flow spectrometer (Applied Photophysics). Samples constituting protein (4 µM), liposomes (1 mM lipids composed of 70% DOPC, 25% DOPS, and 5% dansyl-PE), and CaCl_2_ (250 µM) were allowed to rapidly mix with an equal volume of EGTA (2 mM) at room temperature (23° C). Before mixing, the samples were allowed to equilibrate for 5 min in their respective syringes of the spectrometer. Endogenous protein tryptophan residues were excited at 295 nm and the emission of the dansyl, due to FRET, was monitored via using a 470 nm long-pass filter (KV470, Schott). The reaction volume was set to 120 µl in the stopped-flow instrument. Using Applied Photophysics Pro-Data SX software, the average traces were fitted with either single or double exponential functions and the kinetic values were selected based on the minimum chi-square value of the fitted curve. A few initial data points (within 1 ms) were omitted from the fit to account for the dead-time of the instrument. The experiments were done in biological as well as technical triplicates.

### Nanodisc-black lipid membrane electrophysiology

Reconstitution of three copies of syb2 with three copies of syt1, syt1 5W or syt7 into nanodiscs and t-SNARE SUVs were performed as previously described^42^. In contrast to previous nanodisc-black lipid membrane (ND-BLM) experiments^42^, the results in Fig. 5 used planar lipid bilayers that were formed by drying lipids that were resuspended in water, rather than decane. See Fig. 5a for an illustration of this procedure. This method was developed as DPPS is not soluble in the typical solvent used to form the bilayers, decane, used in classical approaches. Throughout this manuscript, the same planar lipid bilayer formation strategy was used, regardless of lipid mixture, to ensure that samples can be directly compared. Bilayers were formed by resuspending 30 mM of either DOPC/DOPS or DOPC/DPPS in pure water. Five µl of the lipids were mixed with 1 µl t-SNARE reconstituted SUVs (∼ 400 µM lipids, 400 nM protein) and pipetted as a droplet over the aperture in the bilayer cup. The bilayer cup with the lipid droplet is then placed horizontally in a vacuum desiccator for 60 minutes to evaporate the water and form a lipid film across the aperture. As soon as the lipid is dried, the bilayer cup is placed in the bilayer chamber and 25 mM HEPES, 10 mM KCl pH 7.4 buffer is added to the *trans* chamber. To break through the dried lipid, and ‘unplug’ the cup aperture, press firmly on top of the cup to force a small volume of buffer through the aperture, followed by filling the *cis* chamber with 25 mM HEPES, 100 mM KCl. Next, use a fine tipped paint brush, dipped in decane, to re-seal the cup aperture and form the planar lipid bilayer. Note that while DPPS is not soluble in pure decane, when the lipids are in the presence of water, this mixture of polar and non-polar solutions facilitates the formation of a bilayer harboring DPPS. After the bilayer is formed, a *cis* chamber buffer exchange is performed to remove free-floating lipid and t-SNAREs. Subsequent ND-BLM experiments are conducted and analyzed as previously described^42^. ND-BLM experiments described in Supplementary Fig. 12 used conventional BLM methodology via painting lipids in n-decane to form the planar lipid bilayers. These experiments were performed with bilayers composed of DPhPC-DOPS (80:20).

### Data availability

All data presented in this manuscript can be found in the accompanying source data file.

## References

1. Wolfes AC, Dean C. The diversity of synaptotagmin isoforms. Current Opinion in Neurobiology 63, 198–209 (2020).

2. Brose N, Petrenko AG, Sudhof TC, Jahn R. Synaptotagmin: a calcium sensor on the synaptic vesicle surface. Science 256, 1021 (1992).

3. Li C, Ullrich B, Zhang JZ, Anderson RGW, Brose N, Sudhof TC. Ca2+-dependent and Ca2+-independent activities of neural and nonneural synaptotagmins. Nature 375, 594–599 (1995).

4. Bhalla A, Tucker WC, Chapman ER. Synaptotagmin isoforms couple distinct ranges of Ca2+, Ba2+, and Sr2+ concentration to SNARE-mediated membrane fusion. Mol Biol Cell 16, 4755–4764 (2005).

5. Hui E, Bai JH, Wang P, Sugimori M, Llinas RR, Chapman ER. Three distinct kinetic groupings of the synaptotagmin family: Candidate sensors for rapid and delayed exocytosis. Proceedings of the National Academy of Sciences of the United States of America 102, 5210–5214 (2005).

6. Takamori S, et al. Molecular anatomy of a trafficking organelle. Cell 127, 831–846 (2006).

7. Bradberry MM, et al. Rapid and gentle immunopurification of brain synaptic vesicles. J Neurosci 42, (2022).

8. Matthew WD, Tsavaler L, Reichardt LF. Identification of a synaptic vesicle-specific membrane-protein with a wide distribution in neuronal and neurosecretory tissue. Journal of Cell Biology 91, 257–269 (1981).

9. Vevea JD, Chapman ER. Acute disruption of the synaptic vesicle membrane protein synaptotagmin 1 using knockoff in mouse hippocampal neurons. Elife 9, 24 (2020).

10. Littleton JT, Stern M, Perin M, Bellen HJ. Calcium dependence of neurotransmitter release and rate of spontaneous vesicle fusions are altered in Drosophila synaptotagmin mutants. Proc Natl Acad Sci U S A 91, 10888–10892 (1994).

11. Miledi R. Transmitter release induced by injection of calcium-ions into nerve terminals. Proceedings of the Royal Society Series B-Biological Sciences 183, 421–425 (1973).

12. Augustine GJ. How does calcium trigger neurotransmitter release? Current Opinion in Neurobiology 11, 320–326 (2001).

13. Chapman ER. How does synaptotagmin trigger neurotransmitter release? In: Annual Review of Biochemistry). Annual Reviews (2008).

14. Bai H, et al. Different states of synaptotagmin regulate evoked versus spontaneous release. Nature Communications 7, 9 (2016).

15. Geppert M, et al. Synaptotagmin-I - A major Ca^2+^ sensor for transmitter release at a central synapse. Cell 79, 717–727 (1994).

16. Liu HS, Dean C, Arthur CP, Dong M, Chapman ER. Autapses and Networks of Hippocampal Neurons Exhibit Distinct Synaptic Transmission Phenotypes in the Absence of Synaptotagmin I. J Neurosci 29, 7395–7403 (2009).

17. Reist NE, Buchanan J, Li J, DiAntonio A, Buxton EM, Schwarz TL. Morphologically docked synaptic vesicles are reduced in synaptotagmin mutants Drosophila. J Neurosci 18, 7662–7673 (1998).

18. Jorgensen EM, Hartwieg E, Schuske K, Nonet ML, Jin Y, Horvitz HR. Defective recycling of synaptic vesicles in synaptotagmin mutants of Caenorhabditis elegans. Nature 378, 196–199 (1995).

19. Nicholson-Tomishima K, Ryan TA. Kinetic efficiency of endocytosis at mammalian CNS synapses requires synaptotagmin I. Proceedings of the National Academy of Sciences of the United States of America 101, 16648–16652 (2004).

20. Sugita S, et al. Synaptotagmin VII as a plasma membrane Ca2+ sensor in exocytosis. Neuron 30, 459–473 (2001).

21. Vevea JD, Kusick GF, Courtney KC, Chen E, Watanabe S, Chapman ER. Synaptotagmin 7 is targeted to the axonal plasma membrane through γ-secretase processing to promote synaptic vesicle docking in mouse hippocampal neurons. eLife 10, (2021).

22. Wen H, et al. Distinct roles for two synaptotagmin isoforms in synchronous and asynchronous transmitter release at zebrafish neuromuscular junction. Proceedings of the National Academy of Sciences of the United States of America 107, 13906–13911 (2010).

23. Wu Z, Kusick GF, Raychaudhuri S, Itoh K, Chapman ER, Watanabe S. Synaptotagmin 7 docks synaptic vesicles for Doc2α-triggered asynchronous neurotransmitter release. bioRxiv, 2022.2004.2021.489101 (2022).

24. Jackman SL, Turecek J, Belinsky JE, Regehr WG. The calcium sensor synaptotagmin 7 is required for synaptic facilitation. Nature 529, 88-+ (2016).

25. Liu HS, et al. Synaptotagmin 7 functions as a Ca2+-sensor for synaptic vesicle replenishment. eLife 3, 18 (2014).

26. Maximov A, et al. Genetic analysis of synaptotagmin-7 function in synaptic vesicle exocytosis. Proceedings of the National Academy of Sciences of the United States of America 105, 3986–3991 (2008).

27. Yao J, Gaffaney JD, Kwon SE, Chapman ER. Doc2 Is a Ca2+ Sensor Required for Asynchronous Neurotransmitter Release. Cell 147, 666–677 (2011).

28. Martinez I, Chakrabarti S, Hellevik T, Morehead J, Fowler K, Andrews NW. Synaptotagmin VII regulates Ca2+-dependent exocytosis of lysosomes in fibroblasts. Journal of Cell Biology 148, 1141–1149 (2000).

29. Reddy A, Caler EV, Andrews NW. Plasma membrane repair is mediated by Ca2+-regulated exocytosis of lysosomes. Cell 106, 157–169 (2001).

30. Bendahmane M, et al. Synaptotagmin-7 enhances calcium-sensing of chromaffin cell granules and slows discharge of granule cargos. J Neurochem 154, 598–617 (2020).

31. Schonn JS, Maximov A, Lao Y, Sudhof TC, Sorensen JB. Synaptotagmin-1 and-7 are functionally overlapping Ca2+ sensors for exocytosis in adrenal chromaffin cells. Proceedings of the National Academy of Sciences of the United States of America 105, 3998–4003 (2008).

32. Tawfik B, et al. Synaptotagmin-7 places dense-core vesicles at the cell membrane to promote Munc13-2-and Ca2+-dependent priming. Elife 10, (2021).

33. Bai JH, Tucker WC, Chapman ER. PIP2 increases the speed of response of synaptotagmin and steers its membrane-penetration activity toward the plasma membrane. Nat Struct Mol Biol 11, 36–44 (2004).

34. Chapman ER, Davis AF. Direct interaction of a Ca2+-binding loop of synaptotagmin with lipid bilayers. J Biol Chem 273, 13995–14001 (1998).

35. Tran HT, Anderson LH, Knight JD. Membrane-Binding Cooperativity and Coinsertion by C2AB Tandem Domains of Synaptotagmins 1 and 7. Biophysical Journal 116, 1025–1036 (2019).

36. Shahin V, Datta D, Hui E, Henderson RM, Chapman ER, Edwardson JM. Synaptotagmin perturbs the structure of phospholipid bilayers. Biochemistry 47, 2143–2152 (2008).

37. Ward KE, Ropa JP, Adu-Gyamfi E, Stahelin RV. C2 domain membrane penetration by group IVA cytosolic phospholipase A(2) induces membrane curvature changes. Journal of Lipid Research 53, 2656–2666 (2012).

38. Martens S, Kozlov MM, McMahon HT. How synaptotagmin promotes membrane fusion. Science 316, 1205–1208 (2007).

39. Hui E, Johnson CP, Yao J, Dunning FM, Chapman ER. Synaptotagmin-mediated bending of the target membrane is a critical step in Ca(2+)-regulated fusion. Cell 138, 709–721 (2009).

40. Bhalla A, Chicka MC, Tucker WC, Chapman ER. Ca2+-synaptotagmin directly regulates t-SNARE function during reconstituted membrane fusion. Nat Struct Mol Biol 13, 323–330 (2006).

41. Das D, Bao H, Courtney KC, Wu L, Chapman ER. Resolving kinetic intermediates during the regulated assembly and disassembly of fusion pores. Nat Commun 11, 231 (2020).

42. Courtney KC, et al. The complexin C-terminal amphipathic helix stabilizes the fusion pore open state by sculpting membranes. Nat Struct Mol Biol 29, 97-+ (2022).

43. Hui E, Gaffaney JD, Wang Z, Johnson CP, Evans CS, Chapman ER. Mechanism and function of synaptotagmin-mediated membrane apposition. Nat Struct Mol Biol 18, 813–U892 (2011).

44. Martens S, McMahon HT. Mechanisms of membrane fusion: disparate players and common principles. Nat Rev Mol Cell Biol 9, 543–556 (2008).

45. Yao J, Kwon SE, Gaffaney JD, Dunning FM, Chapman ER. Uncoupling the roles of synaptotagmin I during endo- and exocytosis of synaptic vesicles. Nat Neurosci 15, 243–249 (2011).

46. Rhee JS, et al. Augmenting neurotransmitter release by enhancing the apparent Ca2+ affinity of synaptotagmin 1. Proceedings of the National Academy of Sciences of the United States of America 102, 18664–18669 (2005).

47. Mackler JM, Drummond JA, Loewen CA, Robinson IM, Reist NE. The C2B Ca2+-binding motif of synaptotagmin is required for synaptic transmission in vivo. Nature 418, 340–344 (2002).

48. Gaffaney JD, Dunning FM, Wang Z, Hui E, Chapman ER. Synaptotagmin C2B Domain Regulates Ca2+-triggered Fusion in Vitro CRITICAL RESIDUES REVEALED BY SCANNING ALANINE MUTAGENESIS. J Biol Chem 283, 31763–31775 (2008).

49. Shapiro HK, Barchi RL. Alteration of synaptosomal plasma cholesterol content: Membrane physical properties and cation transport proteins. Journal of Neurochemistry 36, 1813–1818 (1981).

50. Lange Y, Swaisgood MH, Ramos BV, Steck TL. Plasma membranes contain half the phospholipid and 90-percent of the cholesterol and sphingomyelin in cultured human fibroblasts. J Biol Chem 264, 3786–3793 (1989).

51. Kiessling V, et al. A molecular mechanism for calcium-mediated synaptotagmin-triggered exocytosis. Nat Struct Mol Biol 25, 911-+ (2018).

52. Bendahmane M, et al. The synaptotagmin C2B domain calcium-binding loops modulate the rate of fusion pore expansion. Mol Biol Cell 29, 834–845 (2018).

53. Li LB, Vorobyov I, Allen TW. The Different Interactions of Lysine and Arginine Side Chains with Lipid Membranes. Journal of Physical Chemistry B 117, 11906–11920 (2013).

54. Brandt DS, Coffman MD, Falke JJ, Knight JD. Hydrophobic Contributions to the Membrane Docking of Synaptotagmin 7 C2A Domain: Mechanistic Contrast between Isoforms 1 and 7. Biochemistry 51, 7654–7664 (2012).

55. Bao H, et al. Dynamics and number of trans-SNARE complexes determine nascent fusion pore properties. Nature 554, 260–263 (2018).

56. Wu Z, Dharan N, McDargh ZA, Thiyagarajan S, O’Shaughnessy B, Karatekin E. The neuronal calcium sensor Synaptotagmin-1 and SNARE proteins cooperate to dilate fusion pores. eLife 10, (2021).

57. Lynch KL, Gerona RRL, Kielar DM, Martens S, McMahon HT, Martin TFJ. Synaptotagmin-1 Utilizes Membrane Bending and SNARE Binding to Drive Fusion Pore Expansion. Mol Biol Cell 19, 5093–5103 (2008).

58. Wu L, Courtney KC, Chapman ER. Cholesterol stabilizes recombinant exocytic fusion pores by altering membrane bending rigidity. Biophys J 120, 1367–1377 (2021).

59. Zick M, Wickner W. Improved reconstitution of yeast vacuole fusion with physiological SNARE concentrations reveals an asymmetric Rab(GTP) requirement. Mol Biol Cell 27, 2590–2597 (2016).

60. Yao J, Kwon SE, Gaffaney JD, Dunning FM, Chapman ER. Uncoupling the roles of synaptotagmin I during endo- and exocytosis of synaptic vesicles. Nat Neurosci 15, 243–249 (2012).

61. Liu HS, Bai H, Xue RH, Takahashi H, Edwardson JM, Chapman ER. Linker mutations reveal the complexity of synaptotagmin 1 action during synaptic transmission. Nat Neurosci 17, 670-+ (2014).

62. Perin MS, Fried VA, Mignery GA, Jahn R, Südhof TC. Phospholipid binding by a synaptic vesicle protein homologous to the regulatory region of protein kinase C. Nature 345, 260–263 (1990).

63. Bai J, Tucker WC, Chapman ER. PIP2 increases the speed of response of synaptotagmin and steers its membrane-penetration activity toward the plasma membrane. Nat Struct Mol Biol 11, 36–44 (2004).

64. Chicka MC, Hui EF, Liu HS, Chapman ER. Synaptotagmin arrests the SNARE complex before triggering fast, efficient membrane fusion in response to Ca(2+). Nat Struct Mol Biol 15, 827–835 (2008).

65. Rao TC, Passmore DR, Peleman AR, Das M, Chapman ER, Anantharam A. Distinct fusion properties of synaptotagmin-1 and synaptotagmin-7 bearing dense core granules. Mol Biol Cell 25, 2416–2427 (2014).

## Methods references

66. Chicka MC, Hui E, Liu H, Chapman ER. Synaptotagmin arrests the SNARE complex before triggering fast, efficient membrane fusion in response to Ca2+. Nat Struct Mol Biol 15, 827–835 (2008).

67. Courtney KC, Vevea JD, Li Y, Wu Z, Zhang Z, Chapman ER. Synaptotagmin 1 oligomerization via the juxtamembrane linker regulates spontaneous and evoked neurotransmitter release. Proceedings of the National Academy of Sciences 118, e2113859118 (2021).

68. Jo S, Kim T, Iyer VG, Im W. Software news and updates - CHARNIM-GUI: A web-based grraphical user interface for CHARMM. Journal of Computational Chemistry 29, 1859–1865 (2008).

69. Klauda JB, et al. Update of the CHARMM All-Atom Additive Force Field for Lipids: Validation on Six Lipid Types. J Phys Chem B 114, 7830–7843 (2010).

70. Abraham MJ, et al. GROMACS: High performance molecular simulations through multi-level parallelism from laptops to supercomputers. SoftwareX 1-2, 19-25 (2015).

70. Abraham MJ, et al. GROMACS: High performance molecular simulations through multi-level parallelism from laptops to supercomputers. SoftwareX 1-2, 19–25 (2015).

